# Metabolism of host lysophosphatidylcholine in *Plasmodium falciparum*-infected erythrocytes

**DOI:** 10.1101/2023.04.17.537066

**Authors:** Jiapeng Liu, Christie Dapper, Michael Klemba

## Abstract

The human malaria parasite *Plasmodium falciparum* requires exogenous fatty acids to support its growth during the pathogenic, asexual erythrocytic stage. Host serum lysophosphatidylcholine (LPC) is a significant fatty acid source, yet the metabolic processes responsible for the liberation of free fatty acids from exogenous LPC are unknown. Using a novel assay for LPC hydrolysis in *P. falciparum*-infected erythrocytes, we have identified small-molecule inhibitors of key *in situ* lysophospholipase activities. Competitive activity-based profiling and generation of a panel of single-to-quadruple knockout parasite lines revealed that two enzymes of the serine hydrolase superfamily, termed exported lipase (XL) 2 and exported lipase homolog (XLH) 4, are the dominant lysophospholipase activities in parasite-infected erythrocytes. The parasite ensures efficient exogenous LPC hydrolysis by directing these two enzymes to distinct locations: XL2 is exported to the erythrocyte, while XLH4 is retained within the parasite. While XL2 and XLH4 were individually dispensable with little effect on LPC hydrolysis *in situ*, loss of both enzymes resulted in a strong reduction in fatty acid scavenging from LPC, hyperproduction of phosphatidylcholine, and an enhanced sensitivity to LPC toxicity. Notably, growth of XL/XLH- deficient parasites was severely impaired when cultured in media containing LPC as the sole exogenous fatty acid source. Furthermore, when XL2 and XLH4 activities were ablated by genetic or pharmacologic means, parasites were unable to proliferate in human serum, a physiologically-relevant fatty acid source, revealing the essentiality of LPC hydrolysis in the host environment and its potential as a target for anti-malarial therapy.

## INTRODUCTION

Malaria counts among the most devastating infectious diseases in the world today. An estimated 247 million cases and 619,000 deaths were attributed to malaria in 2021^1^. Continued development of new anti-malarial strategies will be required to mitigate the effects of eventual drug resistance against current frontline therapies^2^.

During the pathogenic, asexual stage of the human malaria parasite *Plasmodium falciparum* within erythrocytes, high rates of phospholipid synthesis are required to support the expansion of parasite membranes^3^. Parasites also produce the neutral lipids diacylglycerol (DAG) and triacylglycerol (TAG), which are deposited in lipid droplets and mobilized late in the replication cycle^4, 5^. Although the *P. falciparum* genome encodes enzymes for *de novo* fatty acid (FA) synthesis, these are dispensable in the asexual stage^6^. Thus, the intraerythrocytic parasite is dependent on exogenous sources of fatty acids to support its vigorous anabolic activity. Host serum contains two abundant fatty acid sources: free fatty acids and lysophosphatidylcholines (LPC)^7^. Studies with isotope-labeled LPC have demonstrated that LPC is a significant fatty acid source and that LPC-derived fatty acids are readily incorporated into parasite lipids^8, 9^.

Furthermore, LPC can support parasite growth as sole source of exogenous fatty acids^10^. LPC catabolism also provides choline^9, 11^, an important precursor for phosphatidylcholine (PC) biosynthesis. While the metabolites supplied by LPC catabolism support asexual growth, the depletion of this host lipid plays a regulatory role by enhancing commitment to gametocytogenesis^9^.

Although it has long been evident that asexual *P. falciparum* expresses high levels of lysophospholipase activity^12^, the identities of the enzyme(s) that catalyze exogenous LPC hydrolysis are unknown. This is due in part to the large number of possible candidates and the likelihood of functional redundancy: the *P. falciparum* genome encodes 17 members of the serine hydrolase superfamily that are annotated as “lysophospholipase” through either EC number or InterPro domain assignments (data from Plasmodb.org^13^). While two of these, termed PfLPL1 and PfLPL20, have been shown to have lysophospholipase activity *in vitro*^14, 15^, their contributions to the hydrolysis of exogenous LPC have not been examined. Elucidating the repertoire of enzymes responsible for exogenous LPC hydrolysis is a formidable challenge that is unlikely to yield to a piecemeal approach.

To expedite the discovery of key parasite lysophospholipases, we have pursued complementary chemical biology and genetic approaches focused on the serine hydrolase superfamily. We first developed an assay for *in situ* LPC hydrolysis that enabled the identification of inhibitors that blocked the release of fatty acids form LPC (here, *in situ* refers to processes in intact, infected erythrocytes in culture). Candidate lysophospholipases were interrogated by activity-based protein profiling, leading to a serine hydrolase subgroup consisting of four paralogs. Single and multigenic knockout parasite lines were generated to characterize their roles individually and in combination. Two of these four enzymes were revealed to be the source of nearly all LPC-hydrolyzing activity in asexually-replicating parasites. The consequences of the loss of these activities on parasite fitness was investigated, revealing a critical role for LPC hydrolysis for parasite survival in the presence of human serum.

## RESULTS

### Parasite serine hydrolases catalyze LPC hydrolysis in situ

We sought to identify serine hydrolase-directed, small-molecule inhibitors of LPC hydrolysis *in situ* by adapting an intact-cell assay for the incorporation of a fluorescent fatty acid analog, BODIPY™ 500/510 C4, C9 (C4,C9-FA), into parasite neutral lipids^16^. We reasoned that unlabeled fatty acids derived from LPC hydrolysis would compete with C4,C9-FA and thus reduce lipid-associated fluorescence. To test this, C4,C9-FA labeling of the neutral lipids diacylglycerol (DAG) and triacylglycerol (TAG) in cultured parasites was examined in the presence of exogenous LPC (Fig. 1a). LPC 18:1 reduced C4,C9-FA incorporation in a concentration-dependent manner, with 30 and 60 µM suppressing C4,C9-FA incorporation by >90%. LPC 16:0 was less effective, and lyso-platelet activating factor (lyso-PAF), a non-hydrolysable LPC analog, had no effect (Fig. 1a). We therefore conducted *in situ* LPC hydrolysis assays with 30 µM LPC 18:1 (Fig. 1b).

**Figure 1:**
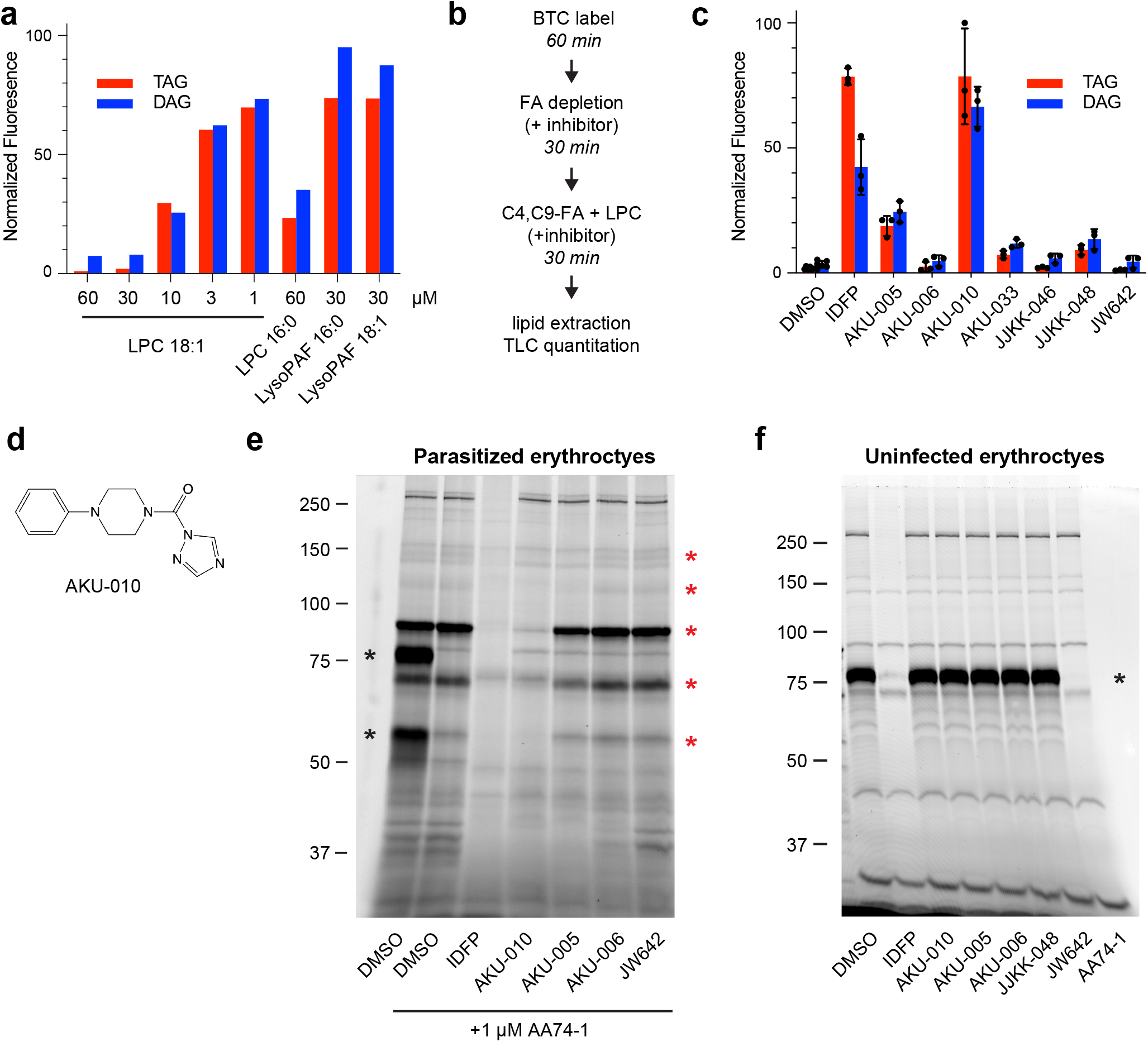
Serine hydrolase inhibitors block LPC hydrolysis *in situ*. (a) Fatty acids derived from LPC hydrolysis reduce C4,C9-FA incorporation into parasite neutral lipids. LPC or the non-hydrolysable LPC analog lyso-PAF were co-incubated with C4,C9-FA and incorporation of the probe into DAG and TAG was quantified and normalized to a no-LPC control (set to 100). (b) C4,C9-FA/LPC competition assay for identifying inhibitors of in situ LPC hydrolysis. (c) Serine hydrolase inhibitors (10 µM) define an inhibition profile for in situ LPC hydrolysis. C4,C9-FA fluorescence volumes were normalized to a no-LPC, no inhibitor control (set to 100). Means and standard deviations are from three independent experiments. (d) Structure of the AKU-010. (e) Inhibition of serine hydrolases in lysates of MACS-enriched, *P. falciparum*-infected erythrocytes as assessed by TAMRA-FP profiling. AA74-1 was included to suppress the strong signal from erythrocyte acylpeptide hydrolase (APEH), which was present at 80 and 55 kDa species (far left lane, black asterisks). Red asterisks indicate species with the inhibition profile AKU-010 > AKU-005 >> AKU-006, JW642. (f) TAMRA-FP profiling of uninfected erythrocytes. Inhibition of erythrocyte serine hydrolases was only observed with IDFP and AA74-1. Black asterisk, APEH. Sizes of markers are indicated in kDa.

We tested a panel of serine hydrolase inhibitors for the ability to block *in situ* LPC hydrolysis as reflected by a gain in C4,C9-FA labeling of DAG and TAG. These inhibitors included: 1) isopropyldodecyl fluorophosphonate (IDFP), a monoacylglycerol isostere and potent lipase inhibitor^17^ that we have previously employed to identify putative *P. falciparum* lipases^18^; 2) JW642, a monoacylglycerol lipase inhibitor^19^ that inhibits an abundant *P. falciparum* serine hydrolase and putative lipase termed “prodrug activating a resistance esterase”, (PfPARE)^18^; and 3) a panel of highly potent, structurally-diverse, monoacylglycerol lipase inhibitors that are based on a piperazine- or piperadine-urea scaffold^20^. All inhibitors act through a competitive-covalent mechanism by reacting with the active site serine residue and were assayed at 10 µM concentration. IDFP treatment resulted in a strong recovery of C4,C9-FA lipid labeling (Fig. 1c), indicating that serine hydrolase-family enzymes are major contributors to LPC hydrolysis *in situ*. Of the other inhibitors, only one, AKU-010^20^, was as effective as IDFP. The structurally-related inhibitors AKU-005 and AKU-006 exhibited modest and no inhibition of LPC hydrolysis, respectively, and JW642 did not inhibit LPC hydrolysis (see Fig. 1d and Supplementary Fig. 1 for compound structures). Thus, compounds AKU-010/005/006/JW642 define an inhibition “signature”, or profile, for the dominant *P. falciparum* lysophospholipase activities.

To identify candidate lysophospholipases, we profiled the inhibitor sensitivities of parasite and host serine hydrolases using activity-based probe TAMRA-fluorophosphonate (TAMRA-FP)^21^. To ensure that all enzymes within the infected erythrocyte were included in this analysis, parasitized erythrocytes (infected RBC, or iRBC) were highly enriched by “magnetic activated cell sorting”, or MACS^22^. Profiling of iRBC lysate revealed multiple enzyme species that matched the *in situ* inhibition profile (Fig. 1e). In contrast, while the host erythrocyte possesses a number of serine hydrolases that are susceptible to IDFP inhibition, none was inhibited by AKU-010 (Fig. 1f), which points to parasite-encoded enzymes being the key catalysts for LPC hydrolysis in infected erythrocytes.

### Two exported serine hydrolases are targets of AKU-010

To gain insight into the enzyme(s) responsible for LPC hydrolysis, we consulted our previous proteomic analysis of serine hydrolase-family lipases in *P. falciparum*^18^. Of the seven putative lipases identified in that study, one stood out as a top candidate: PF3D7_1001600, previously termed “exported lipase 2” (XL2) based on its export to the host erythrocyte cytosol^23^. XL2 has a predicted molecular mass (88.6 kDa) that is consistent with that observed for the major activity in the TAMRA-FP analysis. A paralog of XL2, termed “exported lipase 1” (XL1; PF3D7_1001400), is also exported to the erythrocyte^23^. To determine whether XL1 and XL2 correspond to the AKU-010-sensitive activities, we generated single- and double-knockout parasite lines, targeting the entire coding sequence for deletion using a markerless CRISPR-Cas9 strategy (Supplementary Fig. 2a, b). The ΔXL1 line appeared to have undergone telomere shortening or “healing” (Supplementary Fig. 2c), a phenomenon that can occur when double-stranded DNA breaks are made in subtelomeric regions^24^. The XL2 single knockout and the ΔXL1/2 double knockout contained the expected gene deletion events (Supplementary Figs. 2d, e). All knockout lines grew normally in media containing the serum replacement Albumax I, indicating that neither XL1 or 2 is required under these culture conditions.

Activity-based probe profiling of the knockout lines revealed that XL2 is a highly-expressed enzyme that corresponds to three of the AKU-010-sensitive species (Fig. 2b), the two smaller species likely generated by limited proteolytic cleavage. XL1 is expressed at a much lower level and corresponds to the AKU-010-sensitive activity at ∼110 kDa (Fig. 2a). Both proteins are absent from the ΔXL1/2 line (Supplementary Fig. 2f). To establish that endogenous XL1 and XL2 are exported to the host erythrocyte, MACS-enriched wild-type parasites were fractionated into a supernatant fraction that contains soluble erythrocyte proteins and a pellet fraction that contains intraparasitic proteins (Fig. 2c). Both enzymes were recovered exclusively in the supernatant, confirming efficient export, whereas the intracellular enzyme PfPARE was located in the parasite pellet.

**Figure 2:**
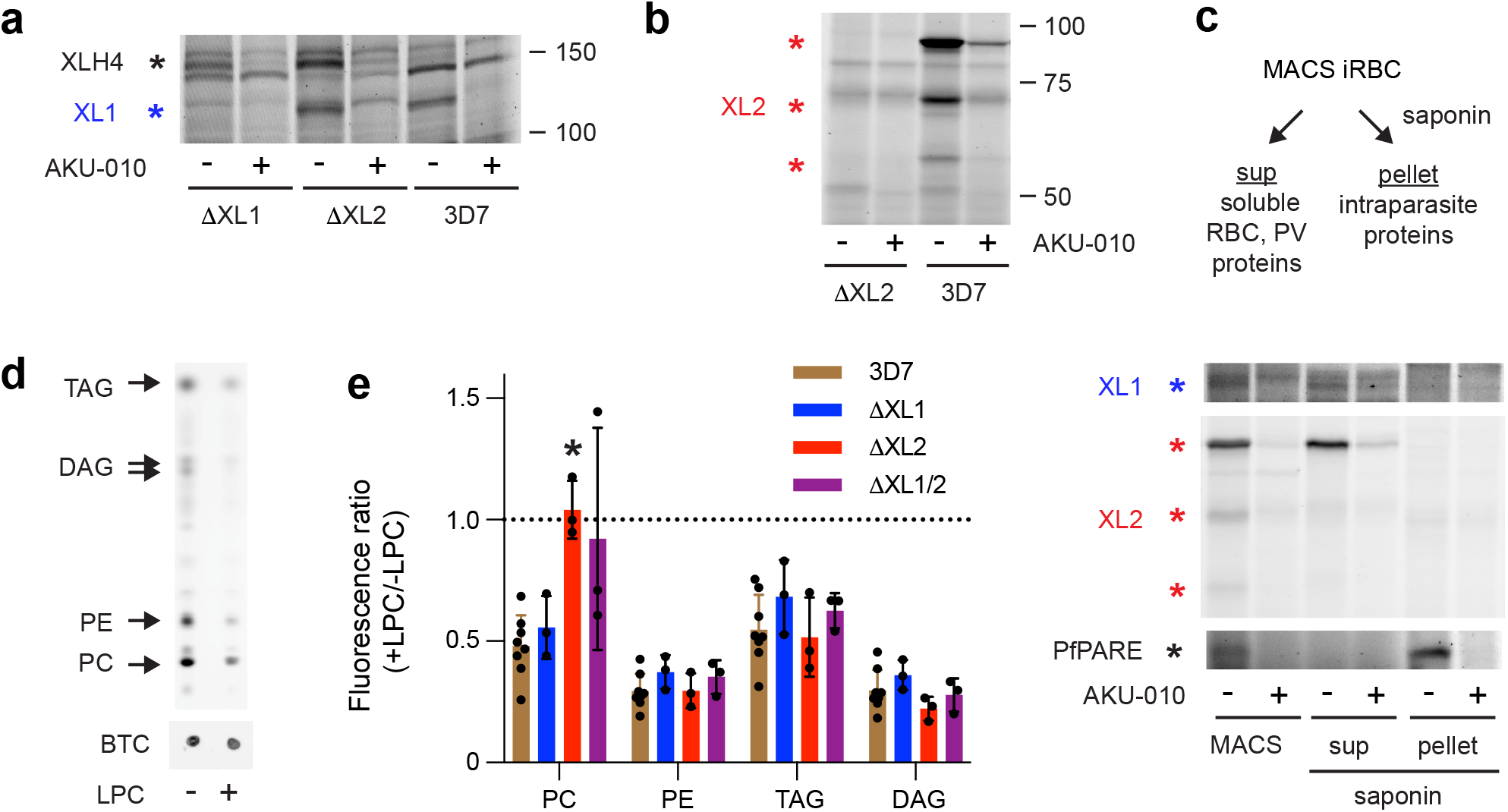
Two exported serine hydrolases are targets of AKU-010. (a) TAMRA-FP profiling of XL1 expression (blue asterisk) in MACS-enriched ΔXL1, ΔXL2 and parental 3D7 iRBCs. An additional, high molecular weight, AKU-010-inhibited species, later identified as XLH4, is indicated with a black asterisk. (b) TAMRA-FP profiling of XL2 expression in MACS-enriched ΔXL2 and 3D7 iRBCs. Three AKU-010-inhibited species corresponding to XL2 are indicated with red asterisks. (c) XL1 and XL2 are exported to the host cell. TAMRA-FP profiling of MACS-enriched 3D7 parasites and of saponin supernatant and pellet fractions. XL1 and XL2 appear predominantly in the saponin supernatant containing soluble erythrocyte proteins, whereas the intracellular serine hydrolase PfPARE appears exclusively in the pellet. (d) LPC reduces incorporation of the fatty acid probe oleate alkyne (OA) into parasite phospholipids (PC, PE) and neutral lipids (DAG, TAG). The two DAG species are 1,2- and 1,3-isomers. BTC, BODIPY-TR-ceramide internal standard. (e) Oleate alkyne/LPC competition profiling of single and double XL1/XL2 knockout lines. Ratios below 1 (dotted line) indicate suppression of OA incorporation in the presence of LPC. Means and standard deviations are from at least three independent experiments. Significance relative to 3D7 was assessed within lipid groups using a two-tailed Welch’s t-test. *, p < 0.05; no asterisk, p ≥ 0.05.

To assess the effects of loss of XL1 and/or 2 on LPC hydrolysis *in situ*, we modified our fatty acid probe-competition assay by replacing C4,C9-FA with a terminal alkyne analog of oleic acid (oleate alkyne, OA), a probe that more closely resembles a physiological fatty acid. Parasites were labeled with 30 µM OA with or without 30 µM LPC 18:1 for 40 minutes. Lipids were extracted from saponin-isolated parasites and alkyne-containing lipids were rendered fluorescent by click addition of 3-azido-7-hydroxycoumarin and resolved by thin-layer chromatography as previously described^25^. For the parental 3D7 line, LPC reduced OA labeling to ∼30-60% of that in its absence (Fig. 2d, e). In ΔXL1, ýXL2 and ΔXL1/2 parasites, no significant changes in OA labeling were observed for PE, DAG or TAG compared to 3D7 (Fig. 2e). In contrast, PC labeling was somewhat elevated in ΔXL2 and ΔXL1/2 lines (this phenomenon is discussed further below). Overall, these studies revealed that XL1- and XL2- deficient parasites retained the ability to efficiently release fatty acids from LPC.

### Intracellular and exported lysophospholipases contribute to efficient LPC hydrolysis

We reasoned that additional AKU-010-sensitive serine hydrolases were responsible for exogenous LPC hydrolysis in ΔXL1/2 parasites. We focused our attention on two enzymes that have been identified as XL1/2 paralogs in the PlasmoDB database: PF3D7_0731800 and PF3D7_1328500. Neither sequence contains a canonical PEXEL host targeting motif^26^ for export to the host erythrocyte (PlasmoDB.org); thus, we refer to these as “exported lipase homolog” (XLH) 3 and 4, respectively.

We first attempted to identify XLH3 in the TAMRA-FP profile of iRBC serine hydrolases by generating a knockout line by marker-free CRISPR/Cas9 deletion (Supplementary Fig. 3a, b). We also generated a yellow fluorescent protein (YFP)-tagged line by single crossover homologous recombination and selection-linked integration^27^ (Supplementary Fig. 4a, b). TAMRA-FP profiling of MACS-enriched iRBCs of the ΔXLH3 and XLH3-YFP parasite lines failed to reveal a labeled species that could be attributed to XLH3, suggesting that the protein is not expressed in asexual stages or is expressed below the detection threshold of our assay. Preliminary OA/LPC competition experiments yielded results essentially identical to those obtained with 3D7; thus, we did not characterize this single-knockout line in detail.

We next generated a parasite line expressing an endogenous XLH4-YFP fusion (Supplementary Fig. 4c). TAMRA-FP profiling revealed a ∼30 kDa increase in molecular mass associated with a ∼150 kDa serine hydrolase that is partially inhibited by 10 µM AKU-010 *in vitro* (Fig. 3a). Live-cell fluorescence microscopy revealed a diffuse cytosolic distribution of XLH4-YFP in the parasite (Fig. 3b), which indicates that XLH4 is not exported to the host cell. To assess the contribution of XLH4 to LPC hydrolysis, we generated parasite lines in which the XLH4 coding sequence was truncated by homologous recombination to render the enzyme non-functional, yielding single (ΔXLH4), double (ΔXL2/XLH4) and triple (ΔXL1/2/XLH4) knockout lines (Supplementary Fig. 3c-e). Finally, we introduced an XLH3 coding sequence deletion on the ΔXL1/2/XLH4 background to produce a parasite line devoid of all four XL/XLH coding sequences (Supplementary Fig. 3f). The resulting ΔXL1/2/XLH3/4 line is referred to as “quadruple knockout”, or QKO. We confirmed the loss of XLH4 in two independently-generated lines (ΔXL2/XLH4 and QKO) by TAMRA-FP profiling (Fig. 3c, Supplementary Fig. 3g).

**Figure 3:**
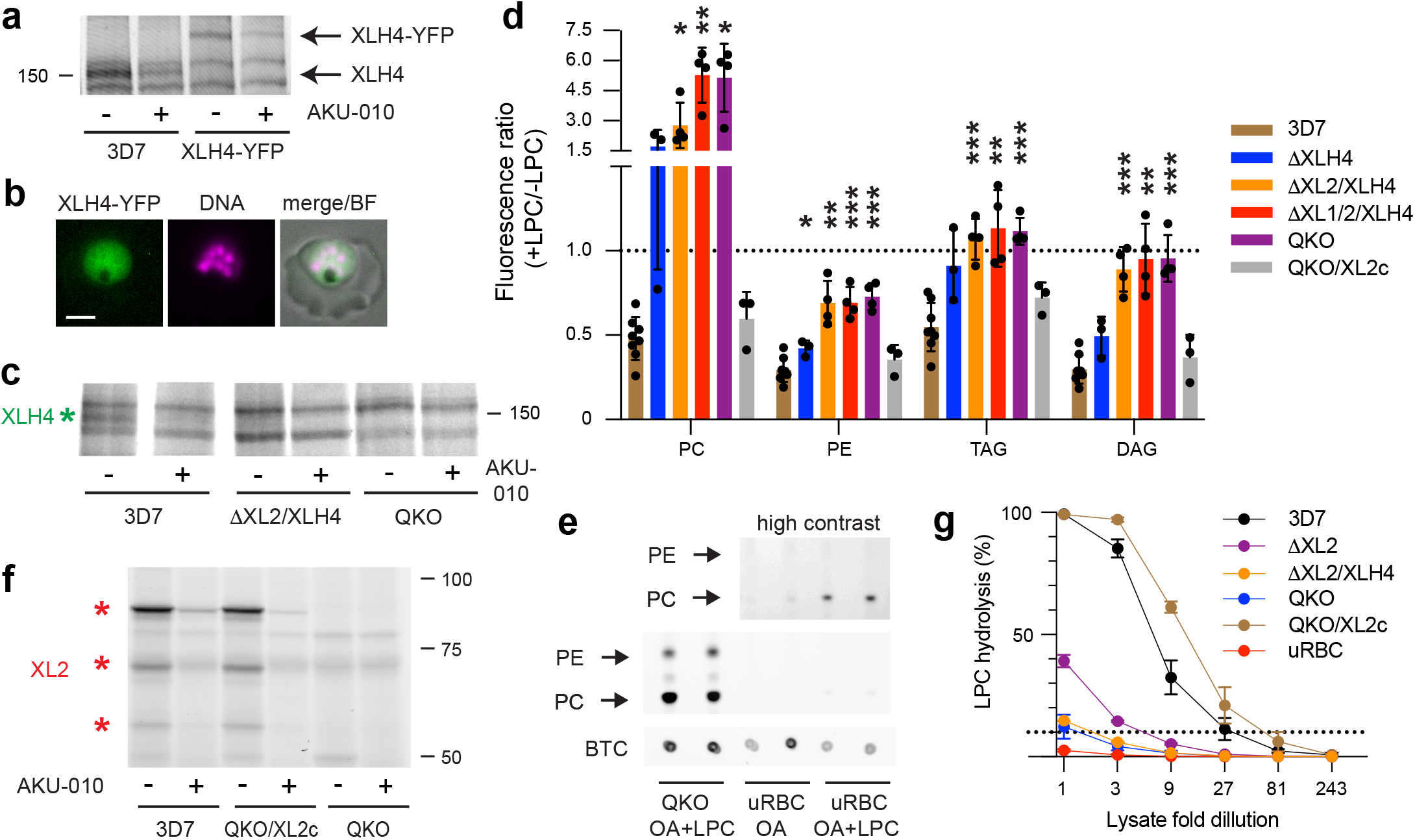
Exported XL2 and intraparasitic XLH4 govern the metabolism of exogenous LPC in asexual parasites. (a) TAMRA-FP profiling of MACS-enriched parental 3D7 and XLH4-YFP-expressing parasites. YFP tagging of XLH4 results in a mass increase of ∼30 kDa. (b) A live parasitized erythrocyte exhibiting intraparasitic XLH4-YFP fluorescence. Hoechst 33342 fluorescence (DNA) is pseudocolored magenta. BF, brightfield. Scale bar, 3 µm. (c) TAMRA-FP profiling demonstrates the loss of XLH4 in MACS-enriched double (ΔXL2/XLH4) and quadruple (QKO) knockout lines. Full gel images are shown in Supplementary Fig. 3g. (d) Oleate alkyne/LPC competition profiling of XLH4 single and XL/XLH multiple knockout lines. A ratio of 1 (dotted line) indicates no competition from LPC hydrolysis. Means and standard deviations are from at least three independent experiments. Significance relative to 3D7 was assessed within lipid groups using a two-tailed Welch’s t-test. *, p < 0.05; **, p < 0.01; ***, p < 0.001; no asterisk, p ≥ 0.05. (e) Enhanced PC synthesis in QKO parasites is not due to erythrocyte acyltransferase activity. Equivalent numbers of MACS-enriched QKO iRBC or uninfected erythrocytes (uRBC) were labeled with OA with or without LPC as indicated, in duplicate. Phospholipids were resolved by TLC. (f) TAMRA-FP profiling of MACS-enriched parasites reveals comparable expression in the complemented QKO parasite (QKO/XL2c) and parental 3D7 lines. (g) *In vitro* lysophospholipase activity in lysates of equivalent numbers of MACS-enriched iRBC or uRBC. Percent of TopFluor LPC hydrolysis is shown for serial three-fold dilutions of lysates. The dotted line indicates 10% substrate hydrolysis. Means and standard deviations are from three independent experiments.

OA/LPC competition assays were conducted to determine the contribution of XLH4 to *in sit*u LPC hydrolysis. While loss of XLH4 alone had a minimal effect, deletion of XLH4 on a ΔXL2 background (ΔXL2/XLH4, ΔXL1/2/XLH4, and QKO lines) was associated with a strong reduction of LPC hydrolysis as reflected by increased OA incorporation into DAG, TAG and PE (Fig. 3d; a +LPC/-LPC ratio of 1 indicates no competition). To our surprise, a ∼10-fold increase in the fluorescence ratio for PC was observed with ΔXL1/2/XLH4 and QKO lines over the parental 3D7 line. This increase was also observed with LPC 16:0, but not when LPC was replaced with LPE, LPS or the non-hydrolysable LPC analog lyso-PAF (all with 18:1 fatty acyl groups; Supplementary Fig. 5). One explanation for this finding is that unhydrolyzed LPC is directly acylated to PC through the activity of an LPC acyltransferase. Because erythrocytes possess LPC acyltransferase activity for the purpose the phospholipid remodeling (*i.e.*, the Lands cycle)^28^, we asked whether PC is efficiently formed from OA and LPC in uninfected erythrocytes. While a very small amount of PC synthesis was observed, this was insignificant compared to that in QKO parasites (Fig. 3e). Thus, the increase in PC synthesis in XL/XLH- deficient parasites cannot be attributed to a host cell acyl transfer pathway.

Finally, we complemented the QKO line with an XL2 expression cassette delivered on a *piggybac* transposon (Supplementary Fig. 6). TAMRA-FP profiling indicated that the complemented parasite line, termed QKO/XL2c, expressed XL2 at a comparable level to that of parental 3D7 parasites (Fig. 3f). XL2 complementation fully restored the capacity of parasites to efficiently hydrolyze LPC *in situ* (Fig. 3d).

### XL2 and XLH4 catalyze LPC hydrolysis

To determine whether XL/XLH enzymes directly contribute to LPC metabolism through catalysis of hydrolysis of the fatty acyl ester (*i.e.*, A1-type lipase activity), we developed an *in vitro* lysophospholipase activity that employs a fluorescent LPC analog, TopFluor-LPC (Supplementary Fig. 7). Lysates of MACS-enriched 3D7, ýXL2, ýXL2/XLH4, QKO and QKO/XL2c parasites, as well as uninfected RBC (uRBC), were generated at equivalent cell densities. Serial 3-fold dilutions were then assayed for hydrolysis of TopFluor-LPC, with substrate and product (TopFluor-oleic acid) resolved by TLC and quantified by fluorescence scanning.

Consistent with a previous report^12^, lysophospholipase (LPL) activity was greatly elevated in 3D7-infected cells over that in uRBC (Fig. 3g). A substantial drop in LPL activity in the ΔXL2 lysate indicates that much of that activity can be attributed to XL2. Interpolating the dilutions required for 10% substrate hydrolysis, we estimate that loss of XL2 resulted in a ∼6- fold reduction in LPL activity. Comparison of ΔXL2 and ΔXL2/XLH4 lysates revealed an additional ∼3-fold drop in LPL activity upon loss of XLH4. No further reduction of LPL activity was observed in QKO parasites, suggesting that neither XL1 nor XLH3 contributed appreciably to hydrolysis of the TopFluor-LPC substrate. The level of LPL activity was slightly above that of uRBC, which suggests that a small amount of residual LPL activity remains in QKO parasites.

XL2 complementation of QKO restored LPL activity to wild-type levels. Thus, XL2 and XL4 are authentic lysophospholipases and constitute the bulk of LPC-hydrolyzing activity in infected erythrocytes.

### XL/XLH-deficient parasites cannot efficiently use exogenous LPC as a source of fatty acids and are hypersensitive to LPC toxicity

We expected that the depletion of LPL activities in QKO parasites would impact their ability to scavenge fatty acids from exogenous LPC. To test this, we washed synchronized ring-stage parasites into media containing two LPC species, 16:0 and 18:1, as sole sources of fatty acids (“2LPC medium”) at concentrations from 5 to 50 µM each. Parasites were also cultured with the two corresponding free fatty acids, palmitate and oleate (30 µM each; “2FFA medium”), and with 0.5% Albumax I, as growth controls. Prior studies have reported that palmitate and oleate are sufficient to support parasite growth over multiple generations^29, 30^; however, we found that only one complete cycle could be sustained in 2LPC or 2FFA medium. Thus, we evaluated parasite proliferation by comparing second-cycle parasitemias, normalizing 2LPC values to those in 2FFA or Albumax-containing media.

Growth of wild-type 3D7 parasites was optimal at LPC concentrations between 15 and 25 µM each, at which the replication efficiency was essentially identical to that in 2FFA medium, but somewhat lower than in Albumax-containing medium (Fig. 4a). Second-cycle parasitemias declined at higher LPC concentrations, presumably due to the adverse effects of elevated LPC concentrations, which have been reported perturb erythrocyte membranes^31, 32^. As expected, the second-cycle parasitemia of QKO parasites in 2LPC medium was substantially diminished at all LPC concentrations compared to that in 2FFA or Albumax (Fig. 4a). Notably, XL2 complementation of the QKO line restored robust growth in 2LPC medium (Fig. 4a).

**Figure 4:**
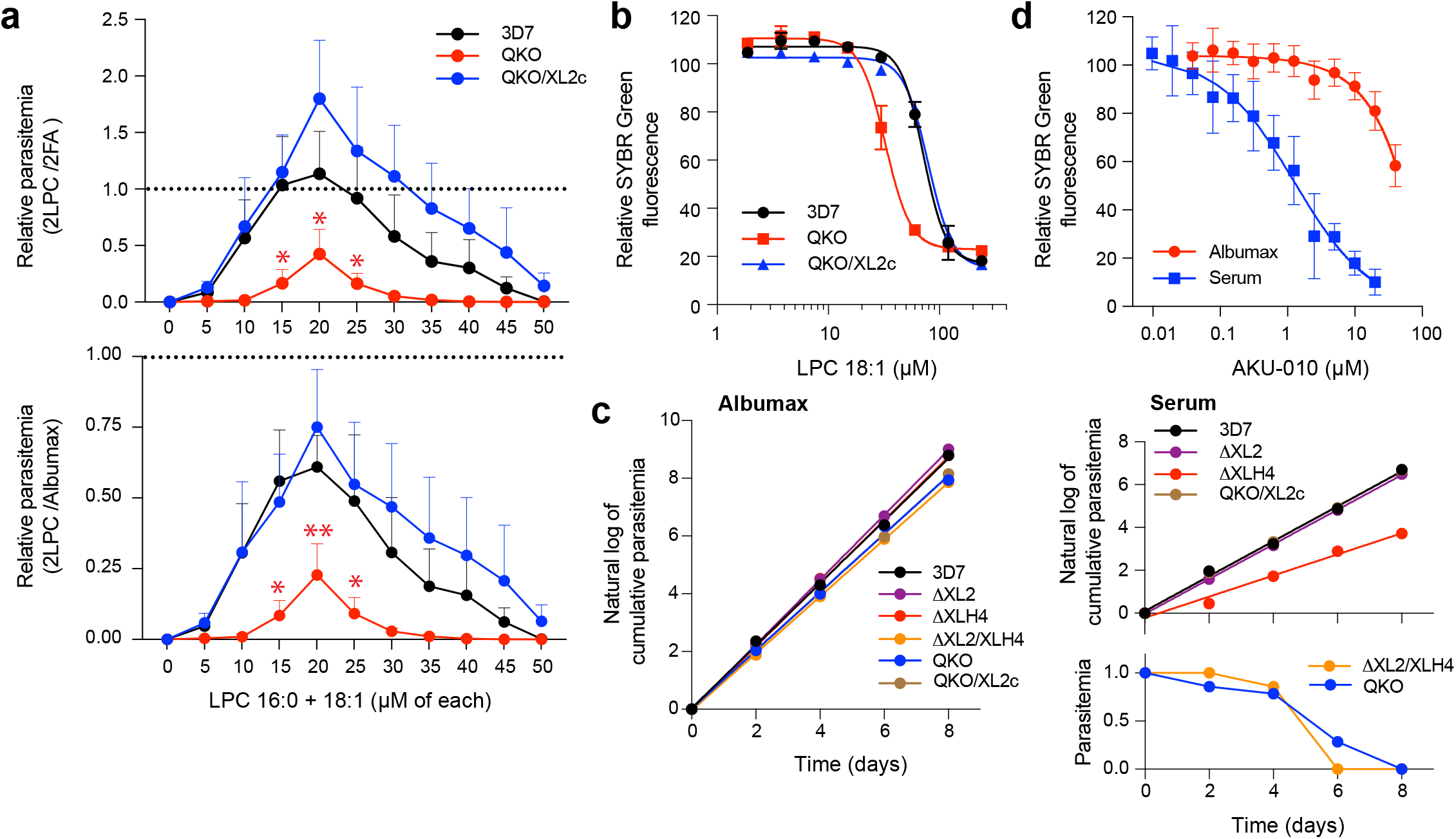
Loss of XL2 and XLH4 activities impairs fatty acid scavenging from LPC, exacerbates LPC toxicity and abrogates growth in serum. (a) QKO parasites have diminished ability to use LPC as a sole source of fatty acids. Parasites were grown for one generation in a minimal-lipid medium containing either two LPC (“2LPC”) or two fatty acid species (“2FA”), or in medium containing Albumax. Parasitemias in 2LPC medium were normalized to those in 2FA medium (upper panel), or in Albumax (lower panel). Means and standard deviations are shown for three independent experiments. Significance relative to 3D7 was assessed for 15-25 uM concentrations using a two-tailed Welch’s t-test. *, p < 0.05; **, p < 0.01. (b) QKO parasites are hypersensitive to LPC toxicity. 3D7, QKO or QKO/XL2c parasites were cultured for 48 h in complete RPMI supplemented with LPC 18:1 (1.9 to 240 µM) and parasite growth was quantified using SYBR Green. Data are from two (3D7, QKO) or one (QKO/XL2c) independent experiments. (c) Parasites lacking XL2 and XLH4 do not proliferate in medium containing human serum. The indicated parasite lines were grown in RPMI medium supplemented with either 0.5% Albumax I or 10 % pooled human serum. Where parasite growth was observed, data were natural-log transformed and fitted by linear regression. One of two independent experiments with similar results is shown. (d) Growth in serum sensitizes wild-type 3D7 parasites to AKU-010. Parasites were incubated with AKU-010 (80 nM - 40 µM for Albumax and 10 nM - 20 µM for serum) for 60 h and parasitemia was quantified on Giemsa-stained smears. Cumulative parasitemia is the parasitemia on day X multiplied by the total dilution up to that point. Means and standard deviations are shown for three independent experiments.

The studies in 2LPC medium suggested that parasite lysophospholipases may fulfill two critical roles: generating fatty acids for lipid synthesis and protecting against LPC toxicity. To evaluate the latter role, we supplemented Albumax-containing medium (which is rich in free fatty acids; see Discussion) with LPC 18:1 at concentrations up to 240 µM and evaluated the growth of 3D7, QKO and QKO/XL2c parasites over one cycle. QKO parasites were substantially more sensitive to LPC toxicity compared to the wild-type 3D7 line (Fig. 4b), with EC_50_ values of 33 ± 3 µM and 72 ± 5 µM, respectively. This sensitivity was reversed by XL2 complementation (Fig. 4b).

We next assessed the capacity of XL/XKH-deficient parasites to proliferate in the presence of a physiologically-relevant, complex source of exogenous fatty acids. Human serum contains high concentrations of free fatty acids and LPC, both in the range of 200-300 µM^7^. Strikingly, QKO parasites were unable to replicate in culture medium containing 10% serum. This phenotype was replicated with ΔXL2/XLH4 parasites but not the single ΔXL2 or ΔXLH4 lines (although the ΔXLH4 line exhibited a reduced growth rate), indicating that robust LPC metabolism is required for parasite growth in the presence of serum. Complementation of the QKO line with XL2 fully restored growth in serum (Fig. 4c). All parasite lines exhibited comparable growth rates in Albumax-containing medium (Fig. 4c), which is likely due to the high free fatty acid and low LPC content of this serum replacement (see Discussion).

Finally, given the inability of LPL-deficient parasites to flourish in the presence of human serum, we asked whether treatment of wild-type 3D7 parasites with the XL/XLH inhibitor AKU- 010 would phenocopy the effects of XL2/XLH4 knockout. This was indeed the case (Fig. 4d). In Albumax-containing medium, parasites were insensitive to AKU-010 concentrations below 10 µM. In contrast, parasite growth in serum-containing medium was inhibited with an EC_50_ value of 1.3 ± 0.4 µM, representing a >10-fold increase in sensitivity to AKU-010, an effect that is likely driven by enhanced LPC toxicity upon inhibition of LPL activity.

## DISCUSSION

We have identified two enzymes, XL2 and XLH4, that are each capable of efficient hydrolysis of exogenous LPC in asexual *P. falciparum*-infected erythrocytes (Fig. 5a). Knockout of either enzyme individually had a negligible effect on the ability of parasites to scavenge fatty acids from LPC; however, deletion of both enzymes greatly reduced parasite lysophospholipase activity *in vitro* and *in situ*, diminished the ability to scavenge fatty acids from LPC, and hypersensitized parasites to LPC toxicity (Fig. 5b). A low level of residual LPL activity in QKO parasites likely accounts for their ability to replicate to a limited extent in 2LPC medium. This LPL activity may derive from two other parasite serine hydrolases that have been shown to have activity *in vitro*; neither of these, however, appears to have as its primary role the hydrolysis of exogenous LPC; rather, they have been associated with vesicular traffic and neutral lipid metabolism^14, 15^.

**Figure 5:**
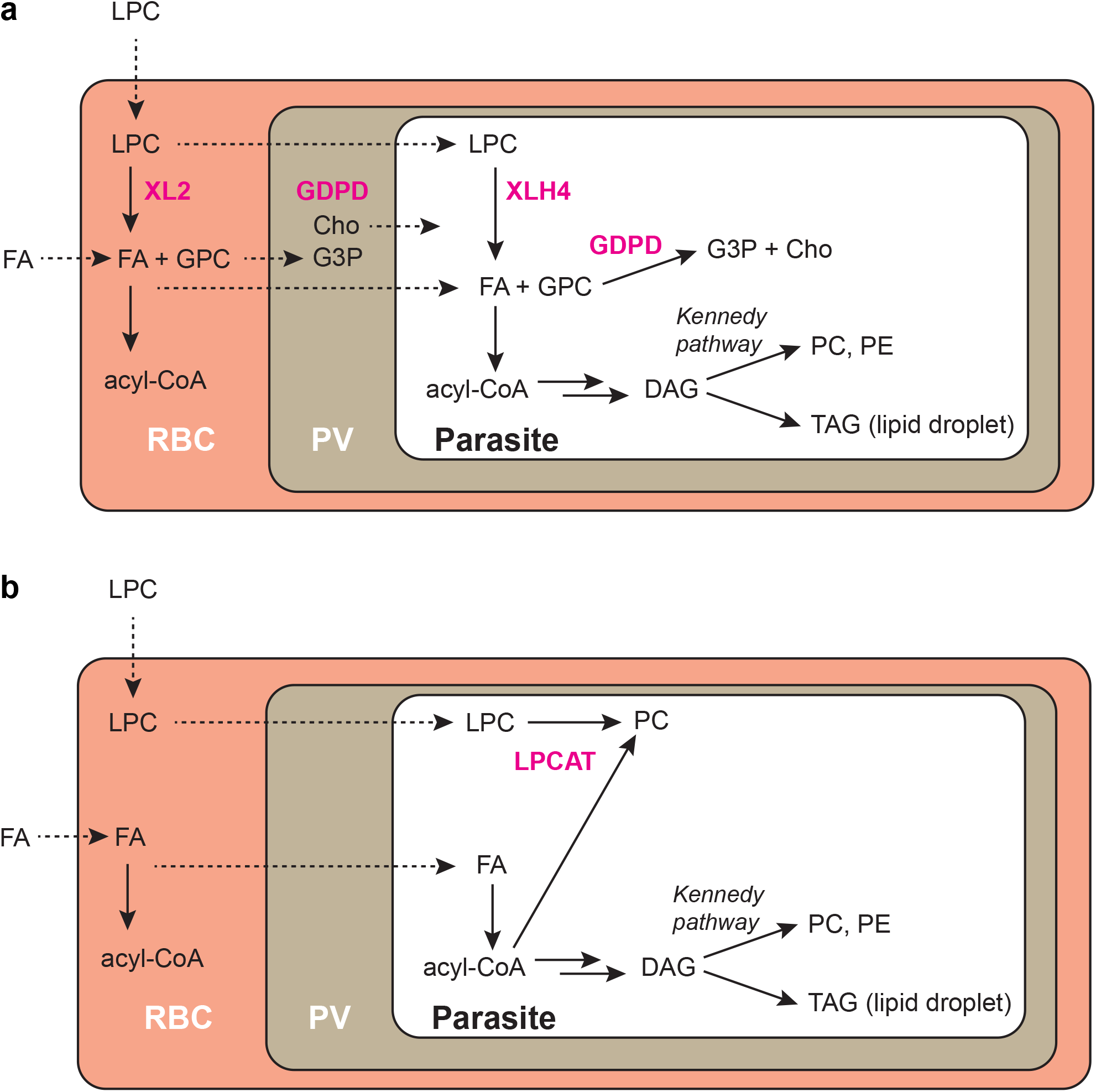
LPC metabolism in *P. falciparum* and metabolic consequences of the loss of XL2/XLH4 activities. (a) LPC is hydrolyzed to fatty acids and glycerophosphocholine (GPC) either in the host erythrocyte by XL2 or within the parasite by XLH4. In both cases, GPC can be hydrolyzed to glycerol-3-phosphate (G3P) and choline (Cho) by glycerophosphocholine phosphodiesterase (GDPD). The resulting fatty acids (FA) serve as precursors for lipid synthesis. Exogenous fatty acids can also be taken up and incorporated into parasite lipids. Solid arrows indicate metabolic reactions with the corresponding enzymes in magenta. Broken arrows indicate transport processes. RBC, red blood cell; PV, parasitophorous vacuole. (b) Loss of XL2 and XLH4 greatly diminishes the rate of hydrolysis of LPC (a low level of hydrolysis is likely occurring but is not depicted). LPC accumulating in the infected erythrocyte may be directly acylated to PC through the activity of an LPC acyltransferase (LPCAT), with acyl-CoA serving as co-substrate.

We propose that XL2 catalyzes the hydrolysis of LPC in the erythrocyte cytosol, yielding fatty acids and glycerophosphocholine (GPC). These fatty acids may serve as substrates for parasite-exported acyl-CoA synthetases^33^ and must also be able to enter the parasite efficiently, given the ability of XL2 complementation of the QKO line to fully restore LPC hydrolysis and growth in 2LPC medium. XL2 is not conserved across *Plasmodium* spp., but rather is restricted to the *Laverania* subgenus consisting of human and ape parasites. This may reflect an adaptation to the mature erythrocyte environment, as these host cells have a limited capacity for lipid metabolism. XLH4 is a cytosolic enzyme that is found in human- and ape-infecting *Plasmodium* species (including the non-Laveranian species *vivax, knowlesi, malariae* and *ovale*) but is absent from rodent parasites (PlasmoDB.org). We speculate that XLH4 catalyzes the hydrolysis of LPC that either escapes hydrolysis in the erythrocyte or that diffuses into the parasite through points of contact between host cell and parasite membranes^34, 35^. Clearly, XLH4 can compensate for the absence of extensive LPC hydrolysis in the host erythrocyte, as indicated by the minimal impact of deleting XL1 and XL2. Fatty acids generated through XLH4 activity can directly enter the parasite acyl-CoA pool and be used for parasite lipid synthesis. While XL2 and XLH4 appear to be functionally redundant in culture, it is possible that they make distinctive contributions to parasite fitness in the host circulation, which contains much higher levels of LPC.

A surprising finding was the ∼10-fold elevation in OA-labeled PC observed in OA/LPC competition assays with parasite lines lacking both XL2 and XLH4, with more modest elevations in the ΔXL2 and ΔXLH4 lines. The most likely explanation for this phenomenon is direct LPC acylation *via* acyl transfer to LPC from acyl-CoA, a ubiquitous reaction in eukaryotes that has not previously been described in P*. falciparum*. According to this model (Fig. 5), intracellular LPC levels are elevated in the absence of XL2 and XLH4, leading to increased flux through an existing LPC acylation pathway. At least two erythrocyte enzymes have LPC acyltransferase (LPCAT) activity: LPCAT1^36^ and peroxiredoxin 6, a trifunctional human enzyme possessing LPCAT activity^37^. These enzymes do not appear to be the putative LPCAT activity observed in XL2/XLH4-deficient parasite lines, however, as the levels of PC production in uninfected erythrocytes were insufficient to account for those in infected erythrocytes. Alternatively, LPCAT activity could be expressed by the parasite; the *P. falciparum* genome encodes a homolog of human LPCAT (PF3D7_0914200). Further investigation will be required to identify the relevant acyltransferase(s) and to evaluate the physiological significance of an LPC acylation pathway in a wild-type genetic background.

Parasite LPC metabolism appears to have a dual role: acquisition of fatty acids and protection from LPC toxicity. All XL/XLH-deficient parasites generated in this study replicated at comparable rates in medium containing Albumax, an enriched bovine serum albumin product that is relatively rich in free fatty acids (FFA) and poor in LPC. Based on literature values for the FFA and LPC content of Albumax^38^, we estimate that our standard formulation of 0.5% Albumax (w/v) in RPMI provided ∼66 µM FFA and ∼11 µM LPC. In contrast, serum contains high levels of both free fatty acids and LPC. A detailed analysis of the human serum metabolome revealed total FFA and LPC concentrations of 220 and 280 µM, respectively (Table 11 in Psychogios *et al*^7^). In medium with 10% (v/v) human serum, the LPC concentration would be ∼28 µM. Thus, we speculate that the elevated LPC concentrations in serum compared to Albumax were responsible for the lack of growth of ΔXL2/XLH4 and QKO parasites in the former. The growth-inhibitory properties of serum could accrue from perturbation to membranes or from dysregulation of PC synthesis in the absence of LPC metabolism. Consistent with this interpretation, wild-type parasites were sensitized to AKU-010 in serum-containing medium.

Together, these findings highlight the importance of XL/XLH-mediated LPC catabolism for parasite proliferation in high-LPC environments such as that found in the host circulation and validate the co-inhibition of XL2 and XLH4 as viable anti-malarial strategy.

## MATERIALS AND METHODS

### Materials

BODIPY™ 500/510 C_4_, C_9_ (C4,C9-FA) and BODIPY-TR-ceramide (BTC) were obtained from ThermoFisher. LPC 18:1 and 16:0, Lyso-PAF 18:1 and 16:0, (*Z*)-octadec-9-en-17-ynoic acid (oleic acid alkyne), TopFluor^®^ LPC, TopFluor^®^ oleic acid, and palmitic and oleic acids were obtained from Avanti Polar Lipids. IDFP and JW642 were purchased from Cayman Chemical. 3- azido-7-hydroxycoumarin was acquired from Abcam. Piperazine-based MAGL inhibitors described in Aaltonen *et al*^20^ were provided by Dr. T. Nevalainen, University of Eastern Finland. The following reagent was obtained through BEI Resources, NIAID, NIH: DSM1, MRA-1161. WR99210 was a gift from D. Jacobus (Jacobus Pharmaceuticals).

### Parasite culture

*P. falciparum* clone 3D7 was routinely cultured in human O^+^ erythrocytes (Interstate Blood Bank) at 2% hematocrit in RPMI 1640 medium supplemented with 0.37 mM hypoxanthine, 11 mM glucose, 27 mM sodium bicarbonate, 10 μg/mL gentamicin and 5 g/L Albumax I (Gibco). Cultures were incubated at 37 °C in a 5% CO_2_ incubator and were synchronized by treatment with 5% (w/v) sorbitol.

### Generation of plasmids and parasite transfection

Oligonucleotides used in plasmid construction and parasite line validation are listed in Supplementary Table 1. All nucleotide sequences were confirmed by DNA sequencing. PCR analyses were conducted on 10 ng of genomic DNA in 15 µL volumes with amplification by Taq polymerase (New England Biolabs). A list of the parasite lines generated in this study is provided in Supplementary Table 2.

Markerless deletion of the entire coding sequences of XL1, XL2 and XLH3 was accomplished by CRISPR/Cas9 editing. Cas9 sgRNAs targeting the respective coding sequences were selected from the database assembled by Ribeiro *et al*^39^ and were cloned into the BtgZ1 site of pUF-Cas9-pre-sgRNA^40^, which contains yeast DHOD as a selectable marker. Homology repair plasmids were constructed using ∼250 nucleotides each of 5’ and 3’ UTR sequences. The 5’ and 3’ homology arms were cloned into *Xho*I/*Sal*I sites and *Eco*RI/*Bgl*II sites of pPM2GT^41^, respectively. 25 µg each of the Cas9/sgRNA plasmid (two Cas9/sgRNA plasmids together were used at 12.5 µg each for XLH3) and the homology repair plasmid (linearized with *Bgl*I) were co-transfected into ring-stage *P. falciparum* by high-capacitance electroporation^42^. After 24 hours, parasites were selected with 2 µM DSM-1 for four days and then grown out without selection. Clonal lines were generated by limiting dilution. Loss of coding sequences was validated by PCR (Supplementary Figs. 2 and 3).

Disruption of the XLH4 coding sequence and generation of C-terminal YFP fusions with XLH3 and 4 were accomplished by homologous recombination and selection-linked integration^27^. The sequence for the yellow fluorescent protein allele Citrine (lacking a stop codon) was inserted into the *Avr*II and *Sal*I sites in pSLI-2×FKBP-GFP^27^, replacing L3-2xFKBP- L4-GFP, to yield pSLI-YFP. An ∼800 bp homology sequence preceded by a stop codon was then inserted into the *Not*I and *Avr*II sites of pSLI-YFP to generate plasmids pSLI-XLH3-YFP, pSLI- XLH4-YFP, and pSLI-XLH4(GD)-YFP. Parasites were transfected by electroporation and after 48 hours were selected with 5 nM WR99210. Resistant parasites were then subjected to selection with 400 µg/mL G418 for up to 12 days to obtain parasites with episomal integration into the targeted genomic locus, which was confirmed by PCR (Supplementary Figs. 3 and 4).

For XL2 complementation, a *piggybac* transposon^43^ carrying a copy of the XL2 coding sequence, with transcription driven by the PfRab7 promoter, was constructed. First, the rab7 5’ UTR (bases -1007 to -1) and the XL2 coding sequence (including a single intron and the stop codon) were amplified from genomic DNA. Both products were merged in an overlapping PCR reaction with oligos 1374/1377 and the resulting fragment was ligated into *Xma*I/*Not*I-digested pDD-mCherry-Rab7-DHOD^44^. QKO parasites were transfected by electroporation and after 48 hours were selected with 1.5 µM DSM-1. Presence of the XL2 coding sequence in drug-resistant parasites was confirmed by PCR (Supplementary Fig. 6).

### C4,C9-FA competition assay for inhibition of LPC hydrolysis in situ

Assays for inhibition of LPC hydrolysis in intact parasitized erythrocytes were conducted using a dual-fluorescent labeling strategy as previously described^16^ and modified as follows. Parasites were pre-labeled with 1 µM BODIPY-TR-ceramide (BTC) in complete RPMI for 1 hour. They were then washed into fatty acid-free (*i.e.*, incomplete) RPMI, aliquoted at 0.5 mL and 8% hematocrit into a 24-well plate, supplemented with 10 µM inhibitor or equivalent volume of DMSO, and incubated for 30 minutes at 37 °C in a CO_2_ incubator. An equal volume of medium containing fatty acid free BSA, C4,C9-FA, LPC18:1 (or 50% ethanol for no-LPC controls), and 10 µM inhibitor (or DMSO for no-inhibitor controls), such that final concentrations were 2 mg/mL BSA, 30 µM C4,C9-FA, 30 µM LPC18:1 and 10 µM inhibitor. After 30 minutes, cultures were transferred to 9 mL of cold 0.03% saponin in PBS and parasites were isolated by centrifugation. Lipids were extracted as previously described^16^, dissolved in chloroform, and spotted on a 10 × 10 cm glass HPTLC Silica Gel 60 plate (Millipore). BTC fluorescence was imaged on a Typhoon RGB imager (Cytiva) using a 633 nm laser and a 670/30 bandpass filter. Neutral lipids were resolved in heptane/diethyl ether/acetic acid at ratio of 40:60:1 and C4,C9-FA fluorescence was imaged using the 532 nm laser and a 570/20 bandpass filter. Fluorescence intensities for DAG, TAG and BTC were quantified using ImageQuantTL (Cytiva). BTC fluorescence intensities were used to normalize DAG and TAG intensities across samples.

### MACS purification and TAMRA-FP analysis

Parasites were enriched from ∼100 mL of culture containing synchronized schizonts (∼36-44 h post-invasion) at ∼10% parasitemia using a SuperMACS™ II Separator and a D Column (Miltenyi Biotec). Numbers of eluted cells were determined with a hemocytometer and parasitemia was calculated from a Giemsa-stained smear. Typical yields were 5 x 10^8^ parasite- infected erythrocytes at 90-95% parasitemia. To generate cell lysates, pellets were resuspended at 5 x 10^8^ parasites/mL in cold PBS supplemented with 5 μM pepstatin A and 10 μM E64. The suspension was sonicated three times for eight seconds using a microtip at 30% maximum power. Hemozoin was removed by centrifugation at 12,000 x*g* at 4 °C. Lysates were aliquoted, snap frozen in liquid nitrogen, and stored at – 80 °C. Lysates of uninfected erythrocytes (uRBC) were generated in the same manner except that the MACS separation was omitted.

For serine hydrolase inhibitor profiling with the activity-based probe TAMRA-FP, 19.4 μL of MACS or uRBC lysate was mixed with 0.2 μL of 1 mM inhibitor (or DMSO for controls), 0.2 µL of 100 µM AA74-1 to selectively inhibit human APEH^45^, and was incubated at 30 °C for 30 minutes. The reaction was added with 0.2 μL of 100 µM TAMRA-FP and incubated at 30 °C for 30 minutes. The reaction was quenched by adding 20 μL of 2X SDS-PAGE loading buffer and incubating at 95 °C for 5 minutes. Labeled proteins were resolved on 8.5 or 10% sodium dodecyl sulfate–polyacrylamide gels and imaged on a Typhoon RGB flatbed scanner using the 532 nm laser and a 570/20 bandpass filter.

### Oleate alkyne competition assay for inhibition of LPC hydrolysis in situ

Synchronized parasite cultures with ∼10% trophozoites were labeled with 1 µM BTC for 1 hour at 37 °C in a CO_2_ incubator and then washed three times with incomplete RPMI. Cultures were resuspended in pre-warmed incomplete RPMI containing 3 mg/mL fatty acid-free BSA, 30 µM oleic acid alkyne and 30 µM LPC 18:1 (or 50% ethanol for no-LPC controls) and incubated for 40 minutes with gentle mixing on an orbital rotator at 37 °C in a CO_2_ incubator. Labeled cultures were harvested and lipids were extracted as described above for C4,C9-FA competition assays. Alkyne-labeled lipids were rendered fluorescent by Cu-catalyzed azide–alkyne cycloaddition of 3-azido-7-hydroxycoumarin as previously described^25^. Solvent was evaporated in a vacuum centrifuge and lipids were redissolved in ∼10 µL chloroform. For TLC separation, 1 µL was spotted on a 10 cm × 10 cm glass HPTLC Silica Gel 60 plate (Millipore). BODIPY-TR-ceramide fluorescence was then recorded as described above for C4,C9-FA competition assays. Phospholipids were developed with 65:25:4:1 chloroform/methanol/water/acetic acid for about 4.5 cm. Plates were air dried and neutral lipids were resolved with 1:1 hexane/ethyl acetate.

Plates were sprayed with 6 mL 4% (v/v) *N*,*N*-diisopropylethylamine in hexane prior to imaging. Fluorescence was imaged on an Azure C400 CCD camera imager using blue LED illumination (472 nm) for excitation and a 513/17 bandpass filter for emission. BTC, neutral and polar lipid fluorescence intensities were quantified using ImageQuantTL software (Cytiva) and lipid intensities were normalized across samples using BTC fluorescence. Competition for OA incorporation was quantified as the ratio of lipid fluorescence in +LPC and -LPC samples

### In vitro LPC hydrolysis assays

Lysates of MACS-purified infected erythrocytes or of uRBC were serially 3-fold diluted with cold PBS. TopFluor LPC was added to 50 µM from a 2.5 mM DMSO stock and reactions were incubated at 30°C for 15 minutes. Reaction products were extracted by transferring 3 µL to 150 µL chloroform and vortexing for 20 seconds. 1 µL of the chloroform layer was spotted on a 10 × 10 cm aluminum HPTLC Silica Gel 60 plate (Millipore). TopFluor LPC and the hydrolysis product TopFluor oleic acid were resolved with 65:25:4:1 chloroform/methanol/water/acetic acid and imaged on a Typhoon RGB scanner using a 488 nm laser and 525/20 bandpass filter. The relative fluorescence intensities of TopFluor LPC and TopFluor oleic acid were assessed by resolving equimolar mixtures using the TLC system described above and quantifying fluorescence intensities. The signal for TopFluor oleic acid was 1.25-fold higher than that for TopFluor LPC; therefore, a correction factor was applied. Percent hydrolysis was then calculated for each sample.

### Growth in medium containing LPC as sole source of fatty acids

Synchronized ring-stage parasite cultures (0-16 h post-invasion) were washed three times with incomplete RPMI and resuspended at 3% parasitemia and 1% hematocrit in 1 mL of RMPI containing 3 mg/mL fatty acid BSA and 0-50 µM each of LPC 16:0 and 18:1 and placed in a 24- well plate. Parallel cultures for normalization were set up with either: 1) RMPI with 3 mg/mL BSA and 30 µM each of palmitic (16:0) and oleic (18:1) acids; and 2) RPMI with 5% Albumax I. Parasites were cultured for 60 hours at 37 °C in a 5% CO_2_ incubator, at which point Giemsa-stained smears were prepared. Parasitemia was calculated from a minimum of 1000 cells.

### Growth in medium containing human serum

Synchronized ring-stage parasite cultures (0-16 h post-invasion) at 2% parasitemia and 2% hematocrit were washed into complete RPMI containing either 0.5% Albumax I or 10% (v/v) pooled, heat-inactivated human serum (Interstate Blood Bank) and cultured at 37 °C in a 5% CO_2_ incubator. Over an eight-day period, parasitemia was counted from Giemsa-stained smears every two days and parasites were subcultured unless they were not thriving, in which case the medium was changed.

### EC_50_ measurements

EC_50_ values for LPC 18:1 and AKU-010 were determined using a SYBR Green I assay as previously described^46^. For LPC, early ring stage parasites (0-8 hours post-invasion) were inoculated into in 96 well plates at 3% parasitemia, 1% hematocrit in complete RPMI containing 0.5% Albumax I. LPC 18:1 (1.9 to 240 µM) or 50% ethanol (final concentration 0.2%) were then added and cultures were incubated in a low-oxygen environment (5% O_2_, 5% CO_2_, and 90% N_2_) at 37°C for 48 hours. No hemolysis was observed under these conditions. For AKU-010, parasites were inoculated in complete RPMI containing 0.5% (w/v) Albumax I or 10% (v/v) pooled human serum. AKU-010 (80 nM - 40 µM for Albumax and 10 nM - 20 µM for serum), DMSO (0.2% or 0.4%), or mefloquine (300 nM; positive control for parasite killing) were then added and cultures were incubated in 5% CO_2_ incubator for 60 hours. Cultures were developed with SYBR Green I as previously described^46^ and fluorescence was quantified on a Molecular Devices SpectraMax M5 microplate fluorometer. Fluorescence values were expressed as a fraction of the DMSO control and the EC_50_ values were determined using four-parameter sigmoidal non-linear regression.

## ACKNOWLEDGMENTS

We are grateful to Tapio Nevalainen (University of Eastern Finland) for providing the monoacylglycerol lipase inhibitors described in Aaltonen *et al*^20^ and to Josh Beck (University of Iowa) for the CRISPR/Cas9 plasmids. M. K. discloses support for this work from National Institutes of Health grant AI133136 and from USDA National Institute of Food and Agriculture HATCH project VA-160082.

## AUTHOR CONTRIBUTION STATEMENT

J.L. and M.K. conceived and designed experiments. J.L. and C.D. conducted all experiments. J.L. and M.K. wrote the manuscript.

