## Supplementary Material for "Metabolism of host lysophosphatidylcholine in *Plasmodium falciparum*-infected erythrocytes"

**Contents**:
Supplementary Figures 1-7
Supplementary Tables 1-2

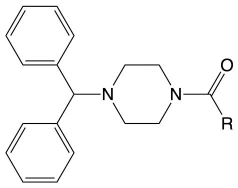


R =

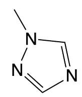


AKU-005

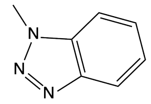


AKU-006

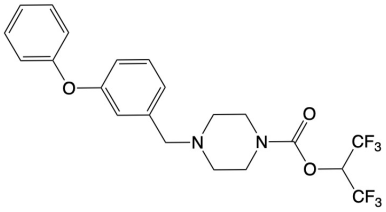


JW642

**Supplementary Figure 1: Structures of inhibitors used for TAMRA-fluorophosphonate profiling of lysophospholipase activities**.

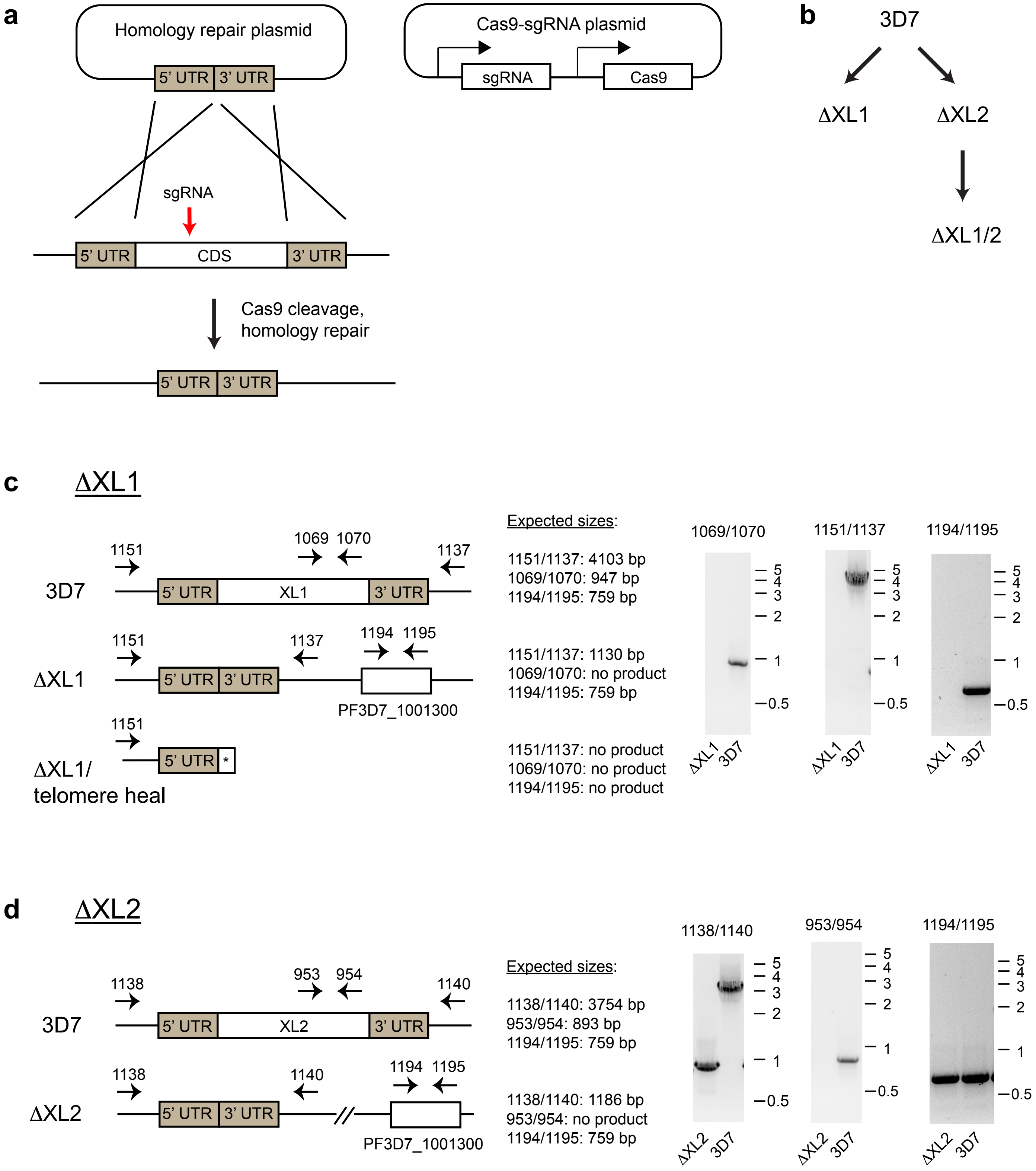


**Supplementary Figure 2:** continued on next page

**e** <u>ΔXL1/2</u>

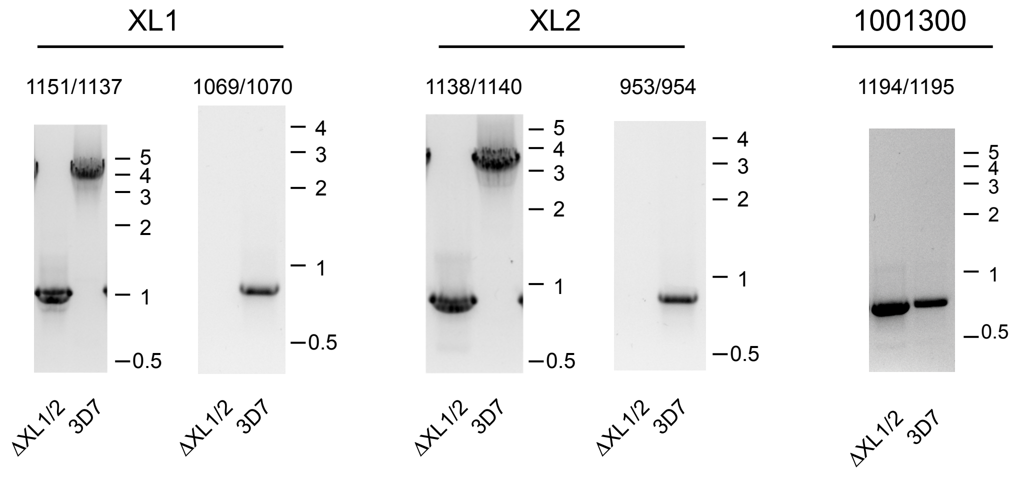


**f** <u>TAMRA-FP profile of ΔXL1/2 line</u>

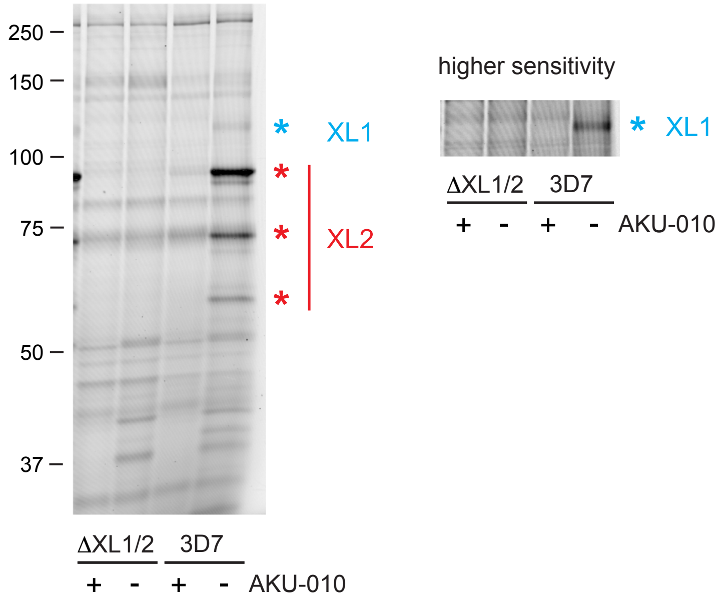


**Supplementary Figure 2: Creation of XL1 and XL2 single and double knockout lines**.
(**a**) CRISPR-Cas9 strategy used for deletion of the entire coding sequences (CDS) of XL1 and XL2. UTR, untranslated region. The sgRNA targets a site in the CDS, shown schematically.
(**b**) Lineage of single and double knockout lines.
**c-e**: PCR analysis of knockout genotypes. Sizes of markers are indicated in kilobases. Introns in XL1, XL2 and PF3D7_1001300 are not shown for clarity.
(**c**) Loss of XL1 occurred through “telomere healing”, evidenced by the absence of PF3D7_1001300.
(**d**) Deletion of the XL2 CDS occurred through the expected homology repair mechanism.
(**e**) Deletion of the XL1 CDS in the ΔXL2 line occurred through the expected homology repair mechanism as indicated by the presence of PF3D7_100130.
(**f**) TAMRA-fluorophosphonate profiling of ΔXL1/2 and 3D7 lines. 1 μM AA74-1 was present to inhibit human APEH. The panel on the right side has been contrast-adjusted to visualize the lighter XL1 band. AKU-010 was added at 10 μM. Sizes of markers are indicated in kilodaltons. Curved lines in the images are Newton’s rings, an interference pattern that occurs during fluorescence scanning when glass plates are closely apposed.

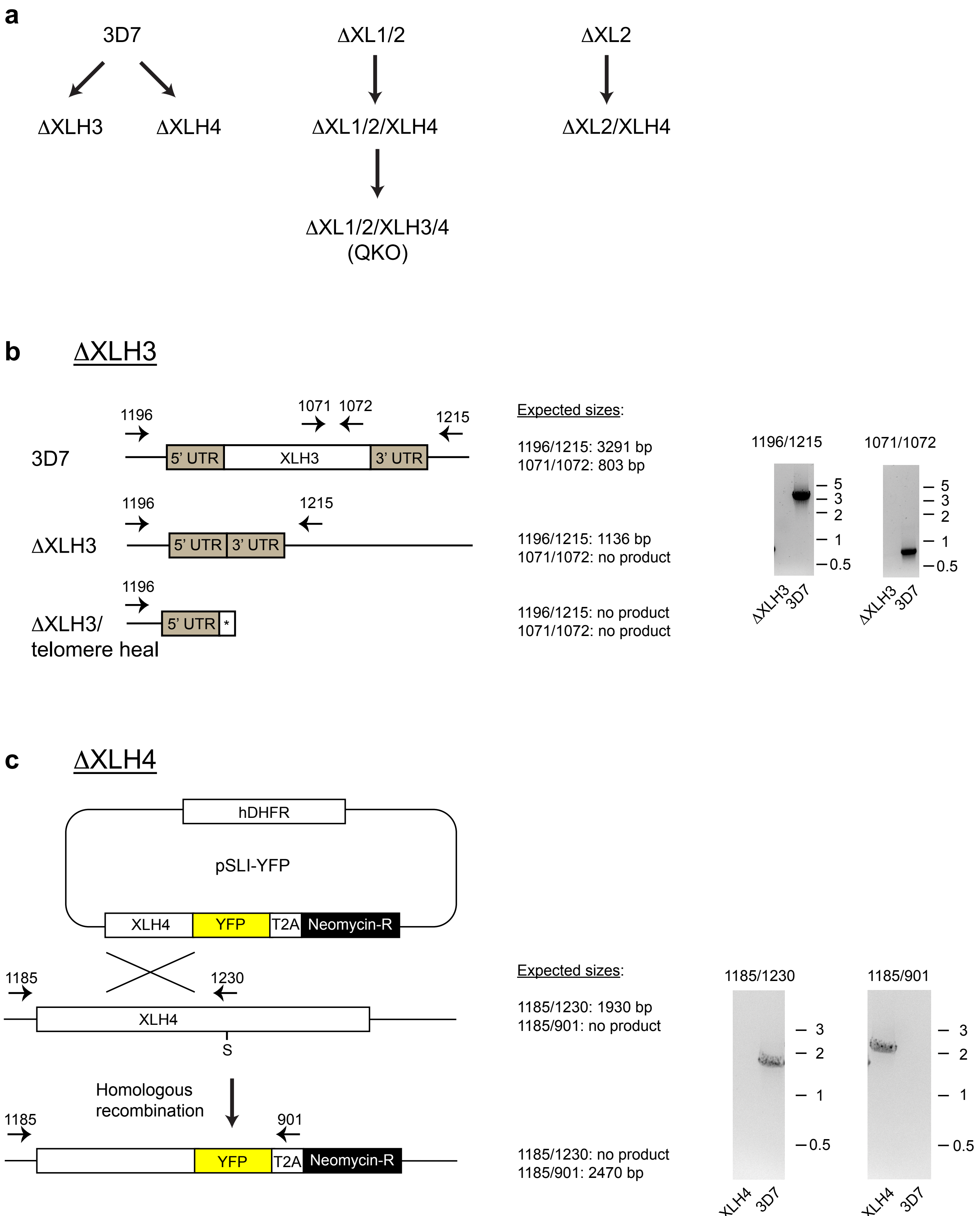


**Supplementary Figure 3:** continued on next page

**d** ΔXL1/2/XLH4

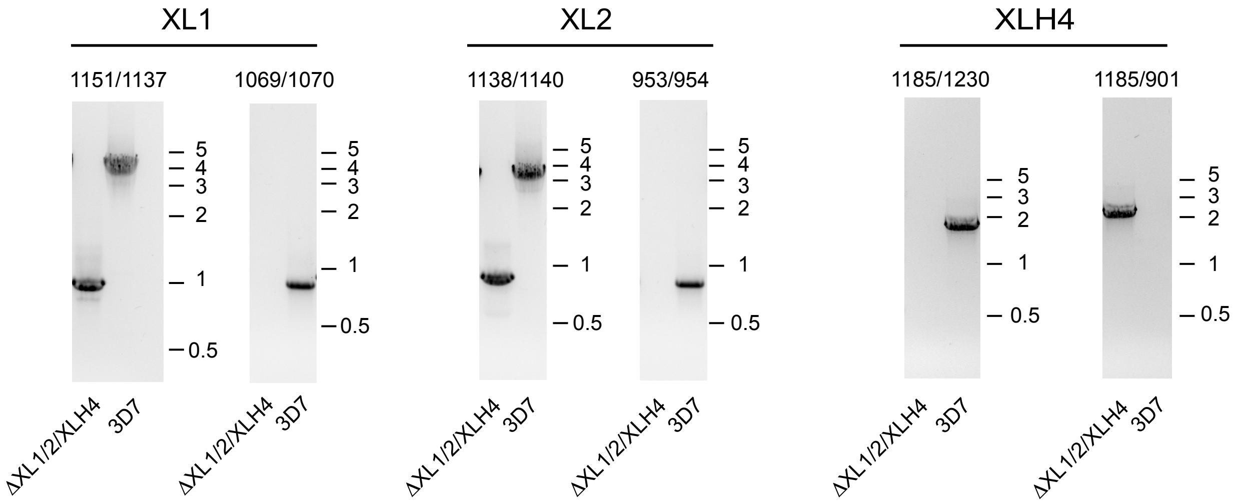


**e** ΔXL2/XLH4

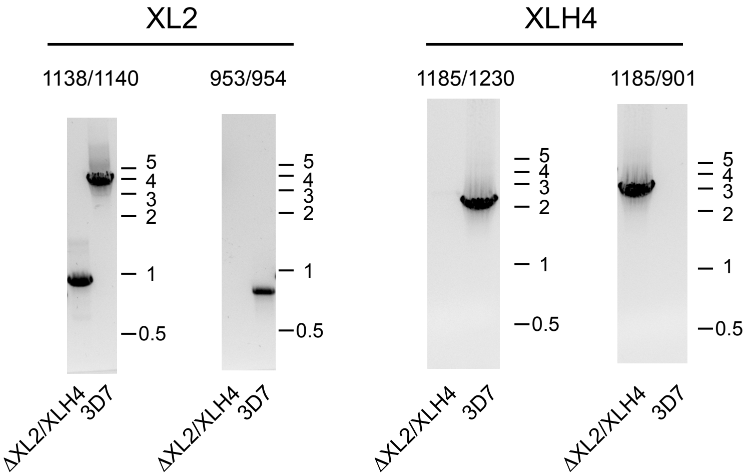


**f** ΔXL1/2/XLH3/4 (QKO)

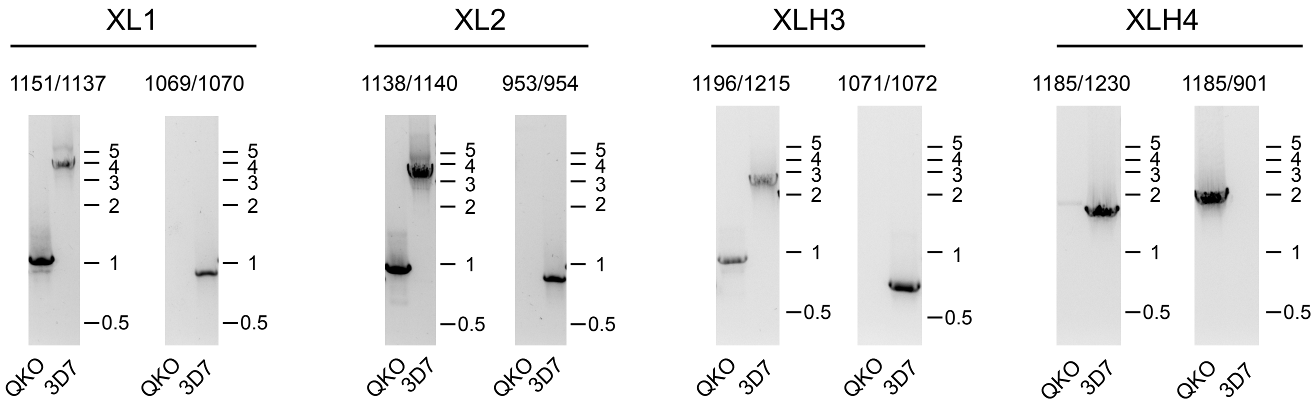


**Supplementary Figure 3:** continued on next page

**g** TAMRA-FP profiles of ΔXL2/XLH4 and QKO lines

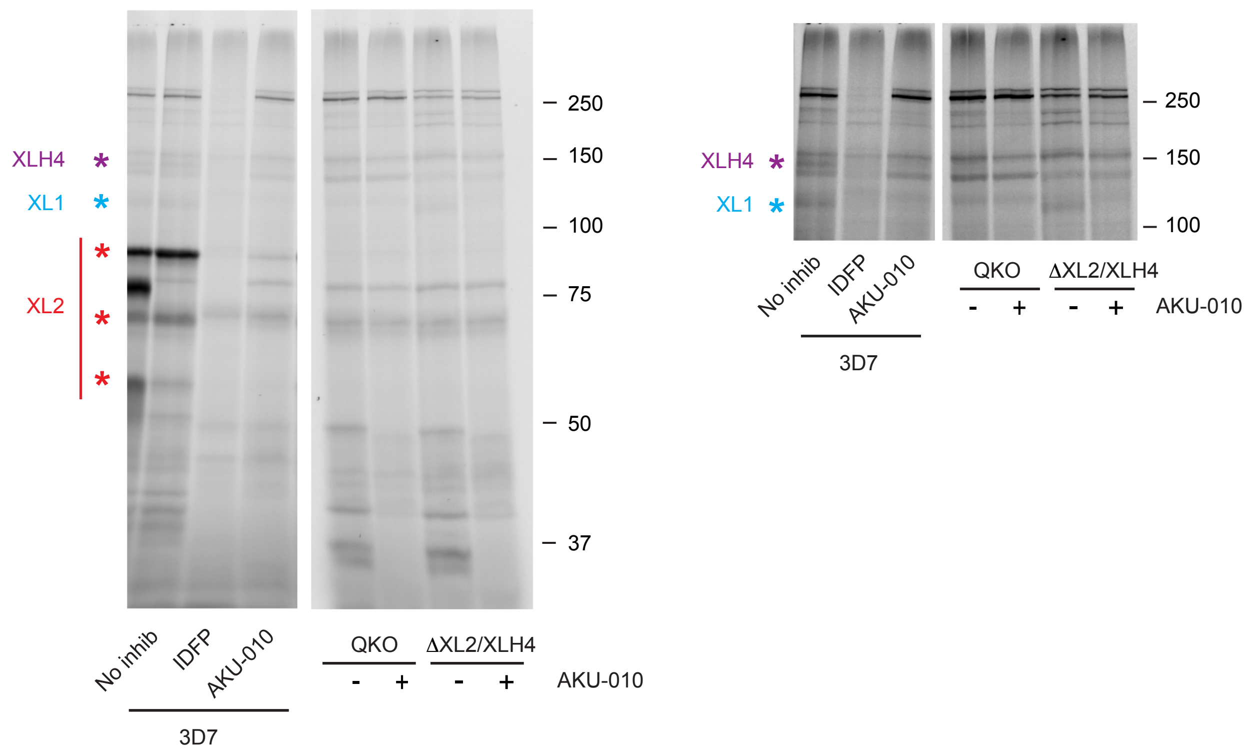


**Supplementary Figure 3: Generation of XLH3 and 4 knockout lines.**
(**a**) Lineages of parasite lines involving disruption of XLH3 and XLH4. QKO, quadruple knockout.
**b-f**: PCR genotyping of knockout lines. See Supplementary Fig. 2 for positions of PCR primers used for XL1 and XL2 genotyping and expected sizes. Positions of markers are indicated in kilobases.
(**b**) Generation of an XLH3 single knockout by markerless CRISPR-Cas9 editing. The Cas9 cleavage was repaired by telomere healing.
(**c**) Disruption of the XLH4 coding sequence through homologous recombination and selection-linked integration. The position of the catalytic serine (S) is indicated. T2A, ribosome skip peptide.
(**d**) Disruption of XLH4 on a ΔXL1/2 background.
(**e**) Disruption of XLH4 on a ΔXL2 background.
(**f**) Disruption of XLH3 on a ΔXL1/2/XLH4 background. XLH3 was repaired by recombination with the donor plasmid, not by telomere healing. QKO, quadruple knockout.
(**g**) TAMRA-fluorophosphonate profiling of two independent XLH4-deficient lineages. 1 μM AA74-1 was present to inhibit human APEH. Panels on the right side have been contrast-adjusted to visualize the lighter XL1 and XLH4 bands. IDFP and AKU-010 were added at 10 μM. Sizes of markers are indicated in kilodaltons.

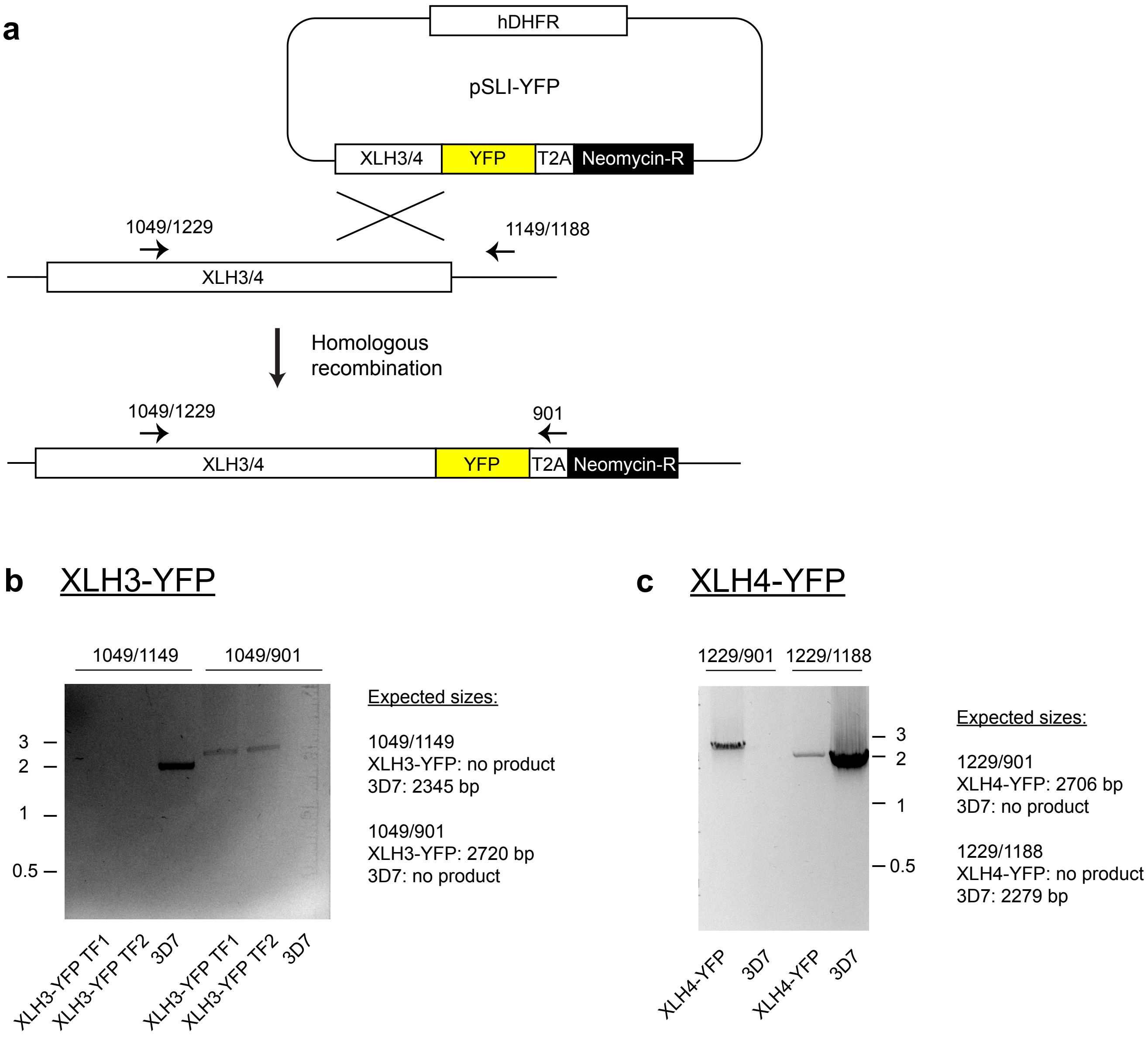


**Supplementary Figure 4: Generation of parasite lines encoding C-terminal YFP fusions with endogenous XLH3 or XLH4.**

(**a**) Strategy for tagging of endogenous loci by homologous recombination and selection-linked integration. PCR primers used for genotyping are shown in the order XLH3/XLH4.

(**b**) Genotyping of XLH3-YFP lines. Two independent transfections (TF1, TF2) were analyzed.

(**c**) Genotyping of an XLH4-YFP line.

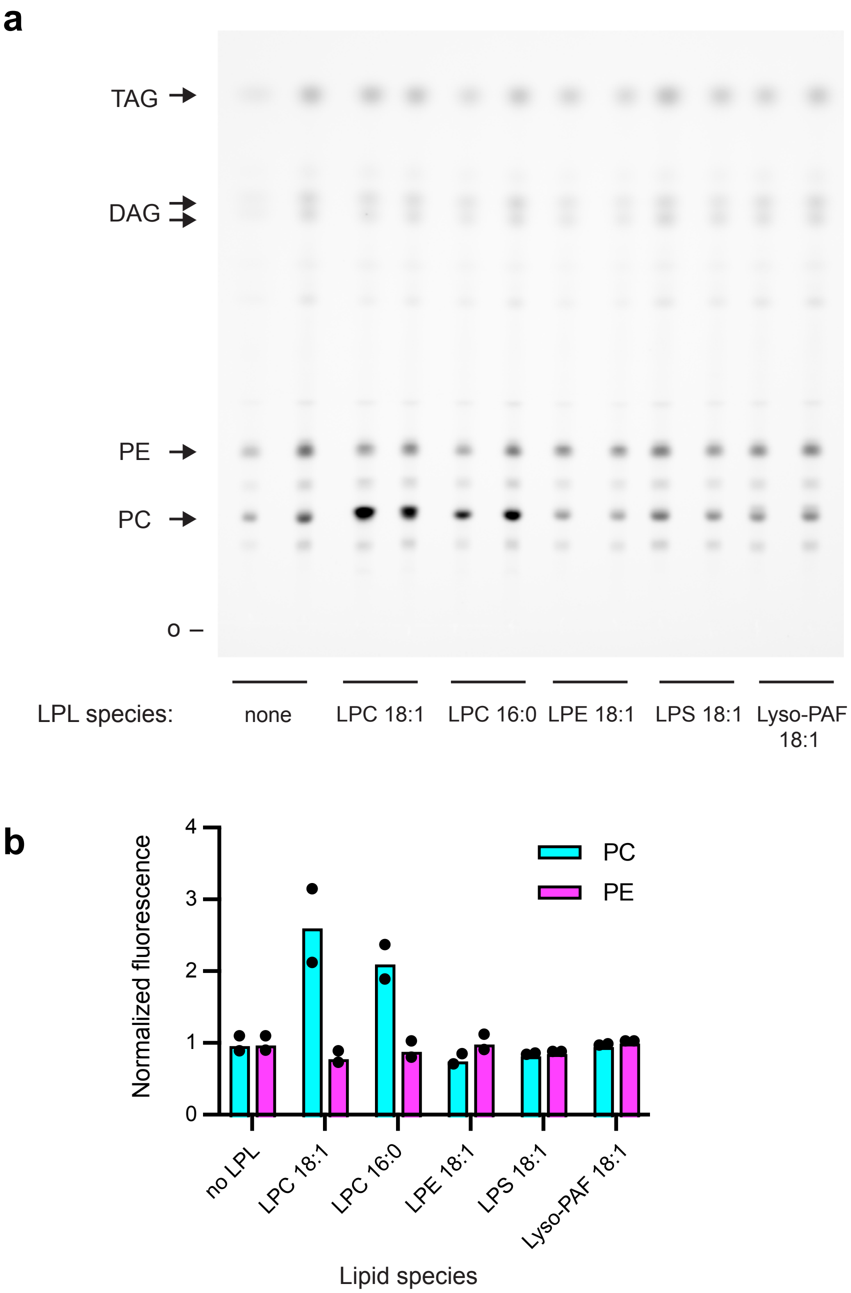


**Supplementary Figure 5: LPC is required for enhanced PC synthesis during oleate alkyne labeling of QKO parasites.**

(**a**) TLC image of oleate alkyne labeling assays conducted with QKO parasites and the indicated lysophospholipids (30 μM). Duplicate cultures were labeled, extracted and spotted for each. o, origin. DAG is resolved into 1,2- and 1,3-isomers. Lyos-PAF, lyso-platelet activating factor, a non-hydrolyzable LPC analog.

(**b**) Quantitation of PC and PE fluorescence intensities. Mean values and individual data points are shown. Fluorescence values were normalized with BTC fluorescence and the mean value for “no LPL” was set to 1.

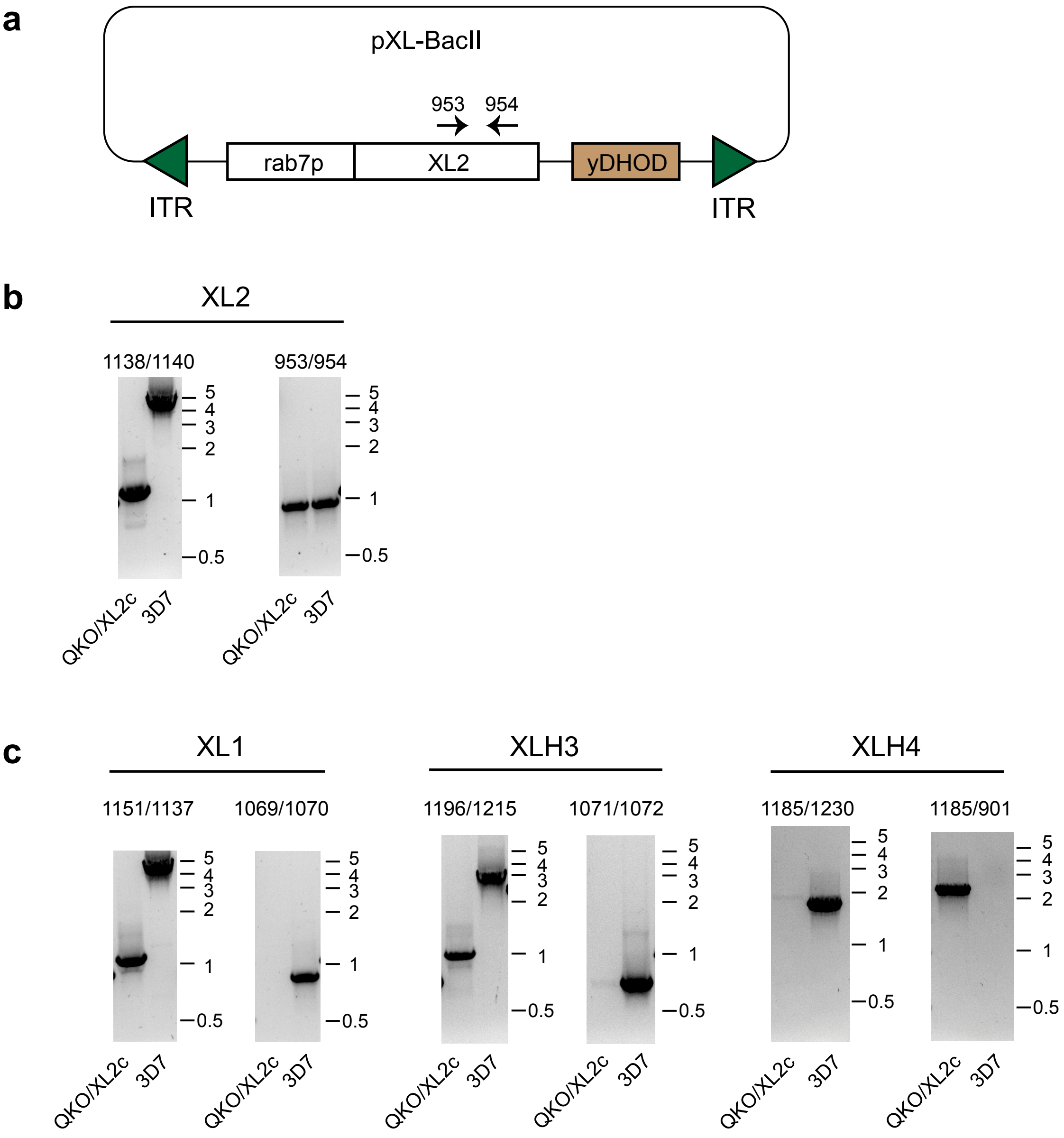


**Supplementary Figure 6: Complementation of the QKO line with XL2**.
(**a**) Design of the *piggybac* transposon-based XL2 expression cassette for complementation of the QKO line (termed QKO/XL2c). The XL2 coding sequence retains the single intron from the genomic sequence (not shown). XL2 expression is driven by the PfRab7 promoter. A yeast dihydroorotate dehydrogenase (yDHOD) cassette confers resistance to DSM-1. ITR, inverted terminal repeat.
(**b**) PCR gentotyping with XL2-specific primers confirms the absence of the genomic coding sequence (1138/1140) and the introduction of a transposon-derived coding sequence (953/954).
(**c**) PCR gentotyping of XL1, XLH3 and XLH4 alleles confirms their knockout status.
See Supplementary Figures 2 and 3 for primer positions and expected sizes of PCR products.

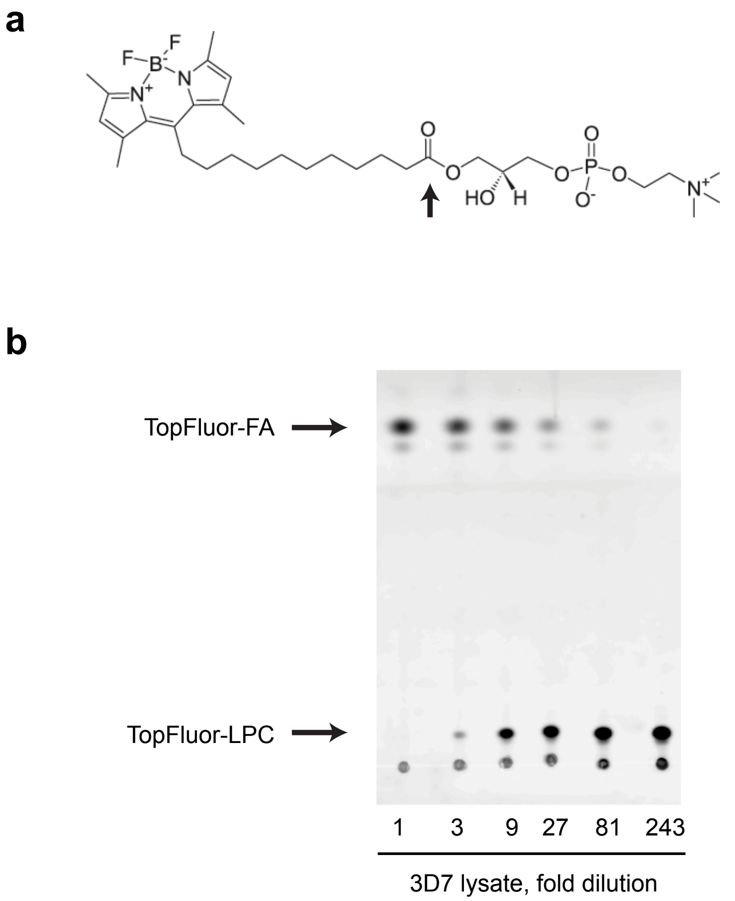


**Supplementary Figure 7: *In vitro* assay for lysophospholipase activity in cell lysates.**
(**a**) Structure of the TopFluor-LPC substrate. The ester bond hydrolyzed by A1-type lysophospholipases is indicated with an arrow.
(**b**) Resolution of TopFluor-LPC and the TopFluor-fatty acid (FA) hydrolysis product by thin layer chromatography. Serial dilution of a lysate of MACS-enriched 3D7 parasites is shown. Positions of TopFluor-LPC and TopFluor-FA standards are indicated with arrows.

**Supplementary Table 1:** Oligonucleotides used in this study. Abbreviations: HA, homology arm; SLI, selection-linked integration; YFP, yellow fluorescent protein; CDS, coding sequence; UTR, untranslated region; GD, gene disruption.

| # | Sequence | Use for: | Sequence |
|---|---|---|---|
| 901 | pSLI | Diagnostic PCR | ATCCATCTTGTTCAATCATTGGTCC |
| 930 | YFP | pSLI-YFP | GTACGCCTAGGGAAAATTTATATTTTCAAAGTATGAGTAAAG |
| 931 | YFP | pSLI-YFP | GTACGGTCGACTTTGTATAGTTCATCCATGCCATG |
| 1101 | XL1, 5' UTR | HA1, CRISPR repair | GTGACACTATAGAACTCGAGATGAATATTCATTTATTCTAAACCTGTG |
| 1102 | XL1, 5' UTR | HA1, CRISPR repair | GGGGATCCTCTAGAGTCGACCAAATTATAAATAAAGTGCCTCTTTTCC |
| 1103 | XL1, 3' UTR | HA2, CRISPR repair | CGGGTACCGAGCTCGAATTCGTTGTTACAAATGTGTAACTGTTATAAT |
| 1104 | XL1, 3' UTR | HA2, CRISPR repair | ATAGGGAGACCGGCAGATCTAATATATAACAATATGAATTCTTTTAAAG |
| 1094 | XL1, CDS | sgRNA | TAAGTATATAATATTAGAACATGTCAGTTCACGAAGTTTTAGAGCTAGAA |
| 1095 | XL1, CDS | sgRNA | TTCTAGCTCTAAAACTTCGTGAACTGACATGTTCTAATATTATATACTTA |
| 1069 | XL1, CDS | Diagnostic PCR, wild-type | TGACACTATAGAATACTCGCGGCCGCTAAGGTGGATGTATAGCTTTAAGAAC |
| 1070 | XL1, CDS | Diagnostic PCR, wild-type | TTGAAAATATAAATTTTCCCTAGGTACAAATATATTATTTATCCAGTCAG |
| 1137 | XL1, 3' UTR | Diagnostic PCR, CRISPR-edited | ATCTGATAAGTTCTCATCTTACAC |
| 1151 | XL1, 5' UTR | Diagnostic PCR, CRISPR-edited | TAAAGCAAAATAATTAAAACTTTATAC |
| 1105 | XL2, 5' UTR | HA1, CRISPR repair | GTGACACTATAGAACTCGAGAACATGAATGACACATATAGTTGTCC |
| 1106 | XL2, 5' UTR | HA1, CRISPR repair | GGGGATCCTCTAGAGTCGACGGTGAATGGATTAATATAAAGCAGTC |
| 1107 | XL2, 3' UTR | HA2, CRISPR repair | CGGGTACCGAGCTCGAATTCTTTCATTCCGCTATAAAAAAATGTTATG |
| 1108 | XL2, 3' UTR | HA2, CRISPR repair | ATAGGGAGACCGGCAGATCTGAATAACTTAATTAATAATATGCACTAG |
| 1098 | XL2, CDS | sgRNA | TAAGTATATAATATTTGCAGATGCACTTTTGAGGGGTTTTAGAGCTAGAA |
| 1099 | XL2, CDS | sgRNA | TTCTAGCTCTAAAACCCCTCAAAAGTGCATCTGCAAATATTATATACTTA |
| 953 | XL2, CDS | Diagnostic PCR, wild-type | TGACACTATAGAATACTCGCGGCCGCTAAAAGTCACAGATTAAAAGTTTAGTAC |
| 954 | XL2, CDS | Diagnostic PCR, wild-type | TTGAAAATATAAATTTTCCCTAGGTGGATATATATTATTAAGCCAATTAAC |
| 1138 | XL2, 5' UTR | Diagnostic PCR, CRISPR-edited | AAAATAGTAACGTCTACTATTAAG |
| 1140 | XL2, 3' UTR | Diagnostic PCR, CRISPR-edited | AACAAATGCATATTCATTAATTGCC |
| 1376 | XL2, CDS | XL2 complement | ATTGAAGGAAAATATGCTCAATTTTAGGAGGTGGAG |
| 1377 | XL2, CDS | XL2 complement | ATTATATAACTCGACGCGGCCGCTTATGGATATATATTATTAAGCCAATTA |
| 1146 | XLH3, 5' UTR | HA1, CRISPR repair | CATACGATTTAGAACAAATAATAACAGTAATGAACTTAATAC |

| | | | |
|---|---|---|---|
| 1147 | XLH3, 5’ UTR | HA1, CRISPR repair | GGGGATCCTCTAGAGTCTTTATTCCAT TAAATTTCCTATAC |
| 1148 | XLH3, 3’ UTR | HA2, CRISPR repair | CGGGTACCGAGCTCGCGTGTGCCTATT TTGTTATTAAAAG |
| 1149 | XLH3, 3’ UTR | HA2, CRISPR repair; diagnostic PCR | ATAGGGAGACCGGCAGGTATTTGGCTT TTACTTAATTTGC |
| 1142 | XLH3, CDS | sgRNA 1 | TAAGTATATAATATTTTATCATGGTAG GCCCGAATGTTTTAGAGCTAGAA |
| 1143 | XLH3, CDS | sgRNA 1 | TTCTAGCTCTAAAACATTCGGGCCTAC CATGATAAAATATTATATACTTA |
| 1144 | XLH3, CDS | sgRNA 2 | TAAGTATATAATATTTCATCATCCAGT TTAACCCAGTTTTAGAGCTAGAA |
| 1145 | XLH3, CDS | sgRNA 2 | TTCTAGCTCTAAAACTGGGTTAAACTG GATGATGAAATATTATATACTTA |
| 1071 | XLH3, CDS | Diagnostic PCR, wild-type; pSLI-XLH3-YFP | TGACACTATAGAATACTCGCGGCCGCT AAGGTAGGACTCTAAAATATCCATGG |
| 1072 | XLH3, CDS | Diagnostic PCR, wild-type; pSLI-XLH3-YFP | TTGAAAATATAAATTTTCCCTAGGGCT AAATATATTATTAAGCCAATCAAC |
| 1196 | XLH3, 5’ UTR | Diagnostic PCR, CRISPR-edited | GATACACACATATGTATAATATACG |
| 1215 | XLH3, 3’ UTR | Diagnostic PCR, CRISPR-edited | ATAGGCATTATTCTAAAATAGCAAC |
| 1049 | XLH3, CDS | Diagnostic PCR, wild-type and XLH3-YFP integration | GGATCGAGGGAAGGATTTCAGAATTC GAAAACCTGTATTTTCAGAGCAATGAA AAATTATATAATTCAAGAAAATA |
| 1275 | XLH4, CDS | pSLI-XLH4-GD | TGACACTATAGAATACTCGCGGCCGCT AAAACGAATTGAAAGAGTCGGAAAAG |
| 1276 | XLH4, CDS | pSLI-XLH4-GD | TTGAAAATATAAATTTTCCCTAGGTTC TCCACAATTACAATAATTACAAAC |
| 1185 | XLH4, 5’ UTR | Diagnostic PCR, wild-type and ΔXLH4 integration | GTGACACTATAGAACTCGAGGTATATT GATGATATACATACGAC |
| 1179 | XLH4, CDS | pSLI-XLH4-YFP | TGACACTATAGAATACTCGCGGCCGCT AAGAGCATAACGAAAAATTGGCGAC |
| 1180 | XLH4, CDS | pSLI-XLH4-YFP | TTGAAAATATAAATTTTCCCTAGGATG AAATATATTATTCAACCAATTCAC |
| 1229 | XLH4, CDS | Diagnostic PCR, wild-type and XLH4-YFP integration | GTGATGATAATATGATGGTTGATG |
| 1230 | XLH4, CDS | Diagnostic PCR, XLH4 wild-type | CACCCATAGATAACCCCATAA |
| 1188 | XLH4, 3’ UTR | Diagnostic PCR, wild-type | ATAGGGAGACCGGCAGATCTCCATTTA AATATATACCTAATTTGGG |
| 1194 | Pf3D7_ 1001300, CDS | Diagnostic PCR | GCTTAATTTATCTAATGGAATATTTG |
| 1195 | Pf3D7_ 1001300, CDS | Diagnostic PCR | ACAATATCATTAAGACAATCTATCC |
| 1374 | Rab7, 5’ UTR | Promoter for XL2 complement | TCGACCTCGATATATCCCGGGGAAAA GTATACAATGACGCATGC |
| 1375 | Rab7, 5’ UTR | Promoter for XL2 complement | TAAAATTGAGCATATTTTCCTTCAATA TATTGTTCTTTTT |

**Supplementary Table 2:** *P. falciparum* 3D7 lines generated in this study.

| Genotype | Clone | Drug resistance |
|---|---|---|
| ΔXL1 | 27 | none |
| ΔXL2 | 43 | none |
| ΔXLH3 | 3 | none |
| ΔXLH4 | 3 | hDHFR, NPT |
| ΔXL1/2 | 51–5 | none |
| ΔXL2/XLH4 | NC | hDHFR, NPT |
| ΔXL1/2/XLH4 | 72 | hDHFR, NPT |
| ΔXL1/2/XLH3/4 (QKO) | 7 | hDHFR, NPT |
| ΔXL1/2/XLH3/4 with XL2 complement | NC | hDHFR, NPT, yDHOD |
| XLH3-YFP | NC | hDHFR, NPT |
| XLH4-YFP | NC | hDHFR, NPT |

Abbreviations: NC, not cloned, hDHFR, human dihydrofolate reductase; NPT, neomycin phosphotransferase; yDHOD, yeast dihydroorotate dehydrogenase.
